# Inferring single-cell protein levels and cell cycle behavior in heterogeneous cell populations

**DOI:** 10.1101/2023.08.24.554605

**Authors:** Bram Thijssen, Hendrika A. Segeren, Qingwu Liu, Lodewyk F.A. Wessels, Bart Westendorp

## Abstract

Individual cells in a genetically identical population can show highly variable behavior. Single-cell measurements allow us to study this variability, but the available measurement techniques have limitations: live-cell microscopy is typically restricted to one or a few molecular markers, while techniques that simultaneously measure large numbers of molecular markers are destructive and cannot be used to follow cells over time. To help overcome these limitations, we present here scMeMo (single cell Mechanistic Modeler): a mechanistic modeling framework that can leverage diverse sets of measurements in order to infer unobserved variables in heterogeneous single cells. We used this framework to construct a model describing cell cycle progression in human cells, and show that it can predict the levels of several proteins in individual cells, based on live-cell microscopy measurements of only one marker and information learned from other experiments. The framework incorporates an uncertainty calibration step that makes the posterior distributions robust against partial model misspecification. Our modeling framework can be used to integrate information from separate experiments with diverse readouts, and to infer single cell variables that may be difficult to measure directly.

## Introduction

Individual cells in a genetically identical population often differ from each other in various ways. This heterogeneity can have a major impact on how cells behave in response to stimuli, stress or treatments. One major aspect in which cells differ from each other is in the cell division cycle. Each cell proliferates at its own speed and timing (Chao *et al*, 2019), and a specific cell can be in any of the phases of the cell cycle. This affects, for example, how a cell responds to DNA damage (Shaltiel *et al*, 2015) and anticancer drugs (Bhuyan *et al*, 1972). Another important difference between cells can be the intracellular concentrations of particular biomolecules. Among others, levels of proteins regulating apoptosis can affect whether a cell dies after exposure to an apoptosis inducer (Spencer *et al*, 2009), and distinct patterns of p21 dynamics can affect whether individual cells continue to proliferate or go into a cell cycle arrest (Hsu *et al*, 2019). Another example is our recent observation that variability in E2F transcription factor activity determines whether cells exit the cell cycle after DNA damage (Segeren *et al*, 2020).

Single-cell measurements allow us to study this variability in behavior within populations of cells. However, there are limitations in the molecular features that can be measured in one experiment. Live-cell microscopy makes it possible to follow single cells over time, but usually requires transduction of cells with relevant biosensors or fluorescent tags to obtain molecular measurements. Typically only up to four molecular markers can be measured simultaneously, as done for example in (Bajar *et al*, 2016). Various other techniques allow measurement of many more molecular features in individual cells, such as through single-cell RNA sequencing (Kolodziejczyk *et al*, 2015), mass cytometry (Bandura *et al*, 2009), iterative immunofluorescence (Gut *et al*, 2018) or single-cell sequencing of DNA-labeled antibodies (van Eijl *et al*, 2018). However, these techniques are destructive and hence only cross-sectional measurements can be obtained. For either of these types of measurements, it would be useful to be able to infer unobserved aspects of individual cells: either infer unobserved molecular species in the case of live-cell imaging, or infer time courses in the case of cross-sectional measurements.

Mechanistic computational models are in principle able to make such single-cell inferences. Various approaches for modeling heterogeneous populations of cells have been developed, with different ways of describing intracellular signaling and different ways of accounting for variability between cells (Zechner *et al*, 2014; Loos *et al*, 2018; Dharmarajan *et al*, 2019; Letort *et al*, 2019; Dixit *et al*, 2020; Lambert *et al*, 2021; Browning *et al*, 2022a, 2022b). These publications have focused on fitting models to data in order to attribute cell-to-cell variability to potential sources of heterogeneity, and to describe or predict distributions of molecular features. We wanted to explore if mechanistic simulations of heterogeneous cell populations can also infer aspects of individual cells – such as the level of a particular protein in a particular cell – rather than inferences about the distribution over all cells. In addition, we wanted to make predictions throughout the cell cycle. This requires integration of several sets of measurements, since it is currently not possible to measure all relevant aspects of the cell cycle in the same cell over time.

To perform single-cell inference with multiple sets of information, we set up a Bayesian inference framework combined with mechanistic simulations of heterogeneous cell populations, which we call scMeMo (single cell Mechanistic Modeler). In scMeMo, we simulate populations of cells, where the individual cells in a population can differ from each other, and use Bayesian inference to estimate the model parameters and the extent of variation in those parameters between cells. To make this computationally tractable, we made several simplifying assumptions. In particular, we assumed that variation in only a subset of model parameters is sufficient to represent the dominant sources of variability between cells. Furthermore, we assumed that these variable rate constants follow a parametric distribution across cells. When these assumptions are applicable, the proposed framework has the benefit that it allows transferring information between experiments.

Models of intracellular signaling networks have to be strong simplifications of the underlying biochemistry, since many details of intracellular processes are not known. In addition, many aspects that are known still have to be left out for computational or practical reasons. To create models that are still reliable, we incorporated a posterior calibration step into scMeMo, that makes the posterior distributions (i.e., the conditional probability of model parameters and predictions, based on the measurements) robust to partial model misspecification. Using the resulting framework, we constructed a model of human cell cycle progression. We trained this model on six available sets of protein and transcript measurements, which were obtained with several different experimental techniques, with three different cell lines, and in different laboratories. These measurements constrained several of the parameters on the model. We then asked the model to make predictions of the levels of six cell cycle proteins, based on traces of only a CDK2 biosensor, and found significant correlations between the predicted and observed levels for five of the six proteins. Thus, we demonstrate that mechanistic simulations of heterogeneous cell populations are a viable strategy to make inferences about protein levels and cell cycle behavior of individual cells.

## Results

### Simulating a population of heterogeneous cells

To describe the behavior of individual cells in a heterogeneous cell population, and to use this model to infer information about individual cells, we aimed to create a robust modeling framework which allowed sufficient variation between cells while keeping the computational cost manageable. We opted to use heterogeneous ordinary differential equations (ODEs) for this. Specifically, each cell is represented by one instance of a system of ODEs, where all cells have the same ODE structure, but the dynamic variables and model parameters can differ between the cells. In principle all model parameters could differ between individual cells, but we made the assumption that variability in only some model parameters is important for the question of interest, and that most model parameters can be assumed to be the same in all cells. It then suffices to simulate a limited number of cells to describe the variability in a whole population, making it is feasible to simulate the entire population during parameter inference. An example model is shown in Figure 1A, and the terminology on model parameters, inference variables and the variability between cells is illustrated in Figure 1B.

**Figure 1:**
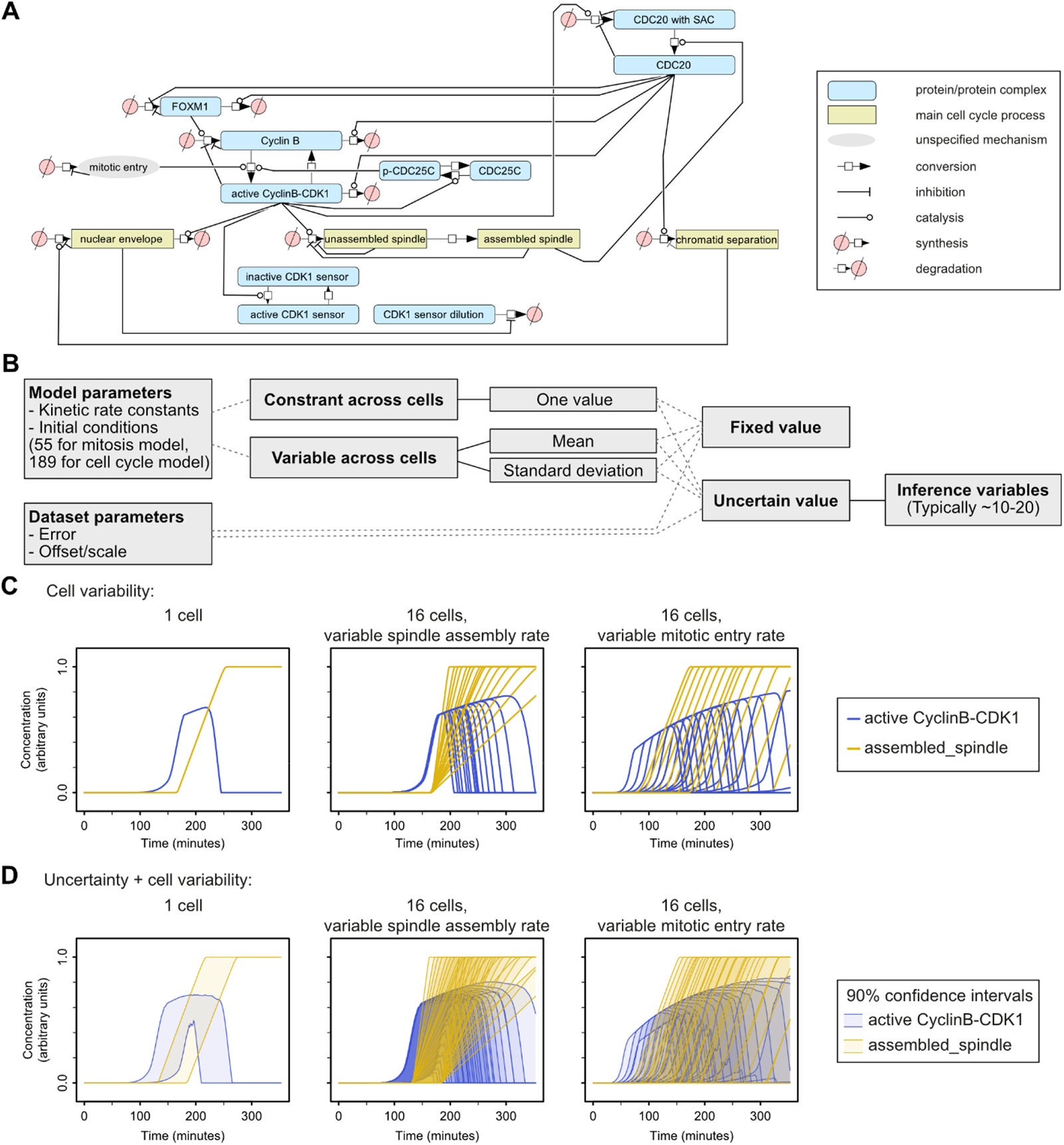
Illustration of the simulation of a population of cells with a simple model of mitosis with scMeMo. **(A)** Structure of the ODE model that is simulated inside every cell, shown in a slightly liberal interpretation of the Systems Biology Graphical Notation format. **(B)** Diagram illustrating the terminology of parameters and variables, and the distinction based on variability across the cell population and whether the values are fixed during inference or not. **(C)** Three simulations illustrating the effect of variability in rate parameters across cells. **(D)** Visualization of the uncertainty. With the Bayesian setup each parameter that is included as inference variable has an uncertainty and this results in a posterior distribution for every dynamic variable, shown here for two dynamic variables. These posterior distributions show only the uncertainty in the dynamic variables in every cell; when fitting data in later figures there is an additional uncertainty on top of the uncertainty in the dynamic variables, describing the residual uncertainty of measurement noise and model error.

Even with a limited number of cells and inferring only a subset of parameters, multivariate parameter inference with such heterogeneous ODEs is still computationally expensive. We therefore incorporated several computational performance optimizations in the model simulation, and furthermore implemented a novel locally adaptive sampling algorithm that efficiently explores the posterior probability distributions. Together, this reduced computation time by a factor fifteen (see the text section Supplementary Result 1 for details). With these optimizations, multivariate parameter inference of 10-20 unknown parameters became feasible, with a parameter inference run typically requiring 24-48 hours on a high performance compute server with 128 cores (with cell populations of several hundred cells and intracellular ODE systems containing several dozen dynamic variables).

### Testing scMeMo with a model of mitosis

To explore whether the scMeMo modeling framework can be used to describe populations of cells, we first focused on progression through mitosis as a test case. We created a simple model that describes several key mitotic events: the production of cyclin B, activation of cyclin B/CDK1 including a positive feedback loop through CDC25C, assembly of the mitotic spindle, and destruction of cyclin B by CDC20 after spindle assembly, followed by chromatid separation (Figure 1A). To illustrate the effect of parameter variability, Figure 1C shows the simulation of a group of cells with this model (using parameter values obtained from model fits described in the next paragraph), where we separately varied two parameters between the cells. When the mitotic spindle assembly rate is varied between cells, the dynamic variable describing the assembled spindle rises faster in some cells than in others. As a result of this variable rate of spindle assembly, the levels of the active cyclin B-CDK1 complex are high for only a brief period of time in some cells, whereas they stay high for a longer period in other cells. On the other hand, when the rate constant describing entry into mitosis is varied but the rate of spindle assembly is held constant, there is variability in when the active cyclin B-CDK1 complex starts to form, while the duration of high cyclin B-CDK1 activity is roughly the same in all cells.

In addition to the cell-to-cell variability described above, there is statistical uncertainty in the trajectories of the dynamic variables. Figure 1D shows this statistical uncertainty by depicting the 90% confidence interval region. Hence, together, there are two sources of variation: a variation between cells due to variability in model parameter values between cells, which we refer to as cell-to-cell variability, and variation for every dynamic variable due to uncertainty in the model parameter values, which we refer to simply as uncertainty.

We fitted the simple mitosis model described above to two sets of measurements: measurements of a CDK1 activity sensor in HeLa cells (Gavet & Pines, 2010), and measurements of eYFP-tagged cyclin B in U2OS cells (Akopyan *et al*, 2014). In fitting these data we allowed three rate constants to vary between cells: the rate controlling when a cell enters mitosis, the rate at which the spindle is assembled, and the rate of FOXM1 synthesis. FOXM1 is a transcription factor responsible for the induction of cyclin B1 expression during late S and G2 phase (Laoukili *et al*, 2007). The reasoning behind choosing only these three rates to vary, is that cell-to-cell variability in these three parameters may be sufficient to describe the main cell-to-cell variability in the population, and that cell-to-cell variability in other model parameters, such as the rate at which CDC25C is phosphorylated, is less important and can be ignored for the present analysis.

Figure 2A shows the fit to the CDK1 FRET sensor measurements. Two sets of measurements were available from the publication of Gavet & Pines; two traces of individual cells (top two panels), as well as the average over six (different) cells aligned at the point of nuclear envelope breakdown (bottom left panel) or at mitotic exit (bottom right panel). The model was fitted to both of these sets of measurements together. The two individual cells clearly differ from each other, with cell 1 having a much shorter duration of high CDK1 activity than cell 2. The model can fit this difference between cells, as shown by the overlay of the measurements and the model fit (Figure 2A). The model used variability in spindle assembly to describe this difference between cells, as indicated by the marginal posterior distribution for the variability of spindle assembly that has a clear peak for values higher than zero (Figure 2B, bottom left panel). Another potential source of cell-to-cell variability, the rate of FOXM1 synthesis, had a much wider marginal posterior probability distribution (Figure 2B, bottom right panel), showing that the extent of cell-to-cell variability in FOXM1 synthesis rate cannot be inferred from these measurements.

**Figure 2:**
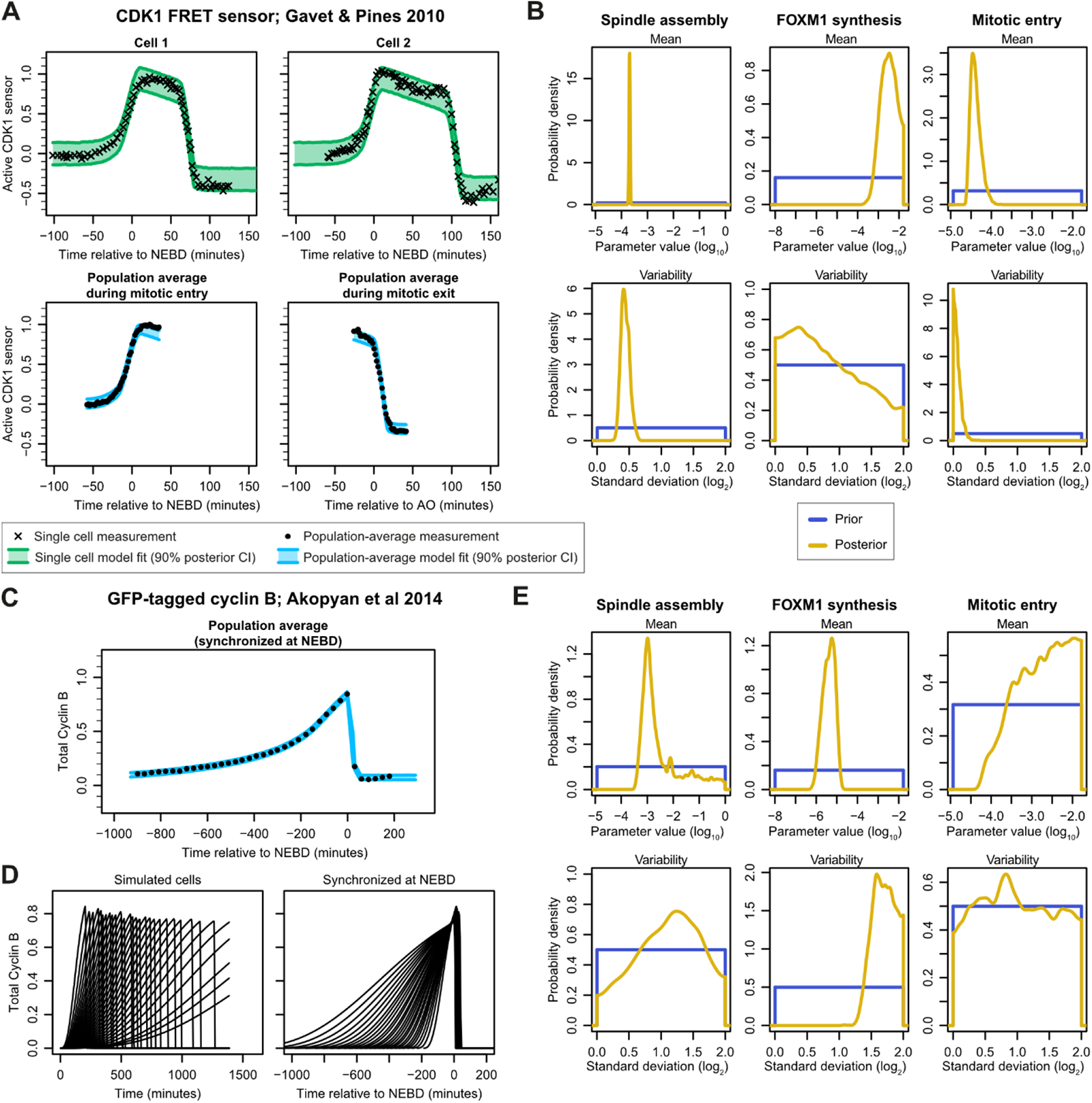
Fit of the simple mitosis model to measurements of a CDK1 sensor and GFP-tagged cyclin B. **(A)** Measurements and 90% posterior confidence interval of the model fit to the CDK1 sensor measurements of Gavet & Pines, for two individual cells and for the average of 6 cells (these are the measurements that were available). In all figures of model fits, crosses and a green shaded area indicates that the measurement and the posterior confidence interval represents an individual cell, and circles and blue shaded area indicates that the measurement and posterior confidence interval represents the average of the entire population. **(B)** Marginal posterior probability distributions of the six inference variables related to the cell-to-cell variability, based on the fit to the measurements from Gavet & Pines. **(C)** Measurements and model fit for the endogenous GFP-tagged cyclin B measurements of Akopyan et al. NEBD = nuclear envelope breakdown. **(D)** Illustration of the alignment of cell trajectories, which produce the steady increase of signal in the averaged population. **(E)** Marginal posterior probability distributions of the six inference variables related to the cell-to-cell variability, based on the fit to the measurements from Akopyan et al.

The fit to the eYFP-tagged cyclin B measurements from the publication of Akopyan et al is shown in Figure 2C. In this case individual traces of cells were not available; only the average of the measurements across multiple cells, after individual traces were aligned at mitotic entry, was provided. However, since we simulate individual cells, we can still fit the model to these averaged data by aligning the simulated single-cell trajectories at mitotic entry and taking the population average at every time point (this was also done for the population average measurements of Gavet & Pines in the previous paragraph). Figure 2D shows simulated trajectories before and after alignment; the left panel illustrates the cyclin B trajectories in the way they are simulated, and the right panel shows the alignment of the individual trajectories at mitotic entry. These aligned trajectories were then averaged across the cells at every time point to fit the averaged measurements. As seen in Figure 2C, the simple mitosis model could describe the averaged trajectory very precisely. In contrast to the measurements from Gavet & Pines, the rate and variability of spindle assembly was less identifiable here (Figure 2E). The FOXM1 synthesis rate and its variability across cells was identifiable however: the model used a clearly nonzero variability of FOXM1 synthesis in order to best describe the fairly long and steady accumulation of cyclin B in the averaged population. Such variability in FOXM1 synthesis fits with multiple factors feeding into FOXM1 expression (Liao *et al*, 2018). Specifically, E2F transcription factors are transcriptional activators of FOXM1 (Millour *et al*, 2011) and E2F-dependent transcription is highly heterogeneous between cells (Segeren *et al*, 2020). Regardless of the underlying mechanism, these results show that an scMeMo model can describe heterogeneity in the process of mitosis in unperturbed human cells.

### Regular Bayesian posteriors are overconfident and require a correction to be robust across datasets

The model used so far is a strong simplification of the complicated mitosis process. Nevertheless, we wondered if this model, using the two limited sets of measurements, can already make accurate predictions. In principle, the cell traces from Gavet & Pines and the averaged traces of Akopyan *et al*, when combined with the assumptions codified in the model, provide important information on the duration of mitosis. We reasoned that measurements of either cyclin B or CDK1 activity over the course of an unperturbed cell cycle reliably predicts the duration of mitotic processes, given the model assumptions on how activation of CDK1 by cyclin B leads to nuclear envelope breakdown and eventual onset of anaphase. To test this, we let the model predict the duration from nuclear envelope breakdown to the onset of anaphase, and compared these predictions to observations of this duration, in the same cell line, in multiple other datasets; either measured by us (denoted Liu et al), or from three publications (Meraldi *et al*, 2004; Lu *et al*, 2013; Zhou *et al*, 2017). The predicted durations were fairly consistent with the observed measurements (Figure 3A). However, two sets of measurements (Meraldi *et al*, and Zhou *et al*) contain a large number of individual cells that progressed from nuclear envelope breakdown to the onset of anaphase more rapidly than the fastest cell in the model prediction. Two explanations for this discrepancy could be that the cell line has diverged between labs, or that experimental conditions differed between experiments. However, we would like model predictions to be robust against such differences, and the posterior uncertainty should therefore have allowed for the apparently faster cells that were seen in the data of Meraldi *et al* and Zhou *et al*.

**Figure 3:**
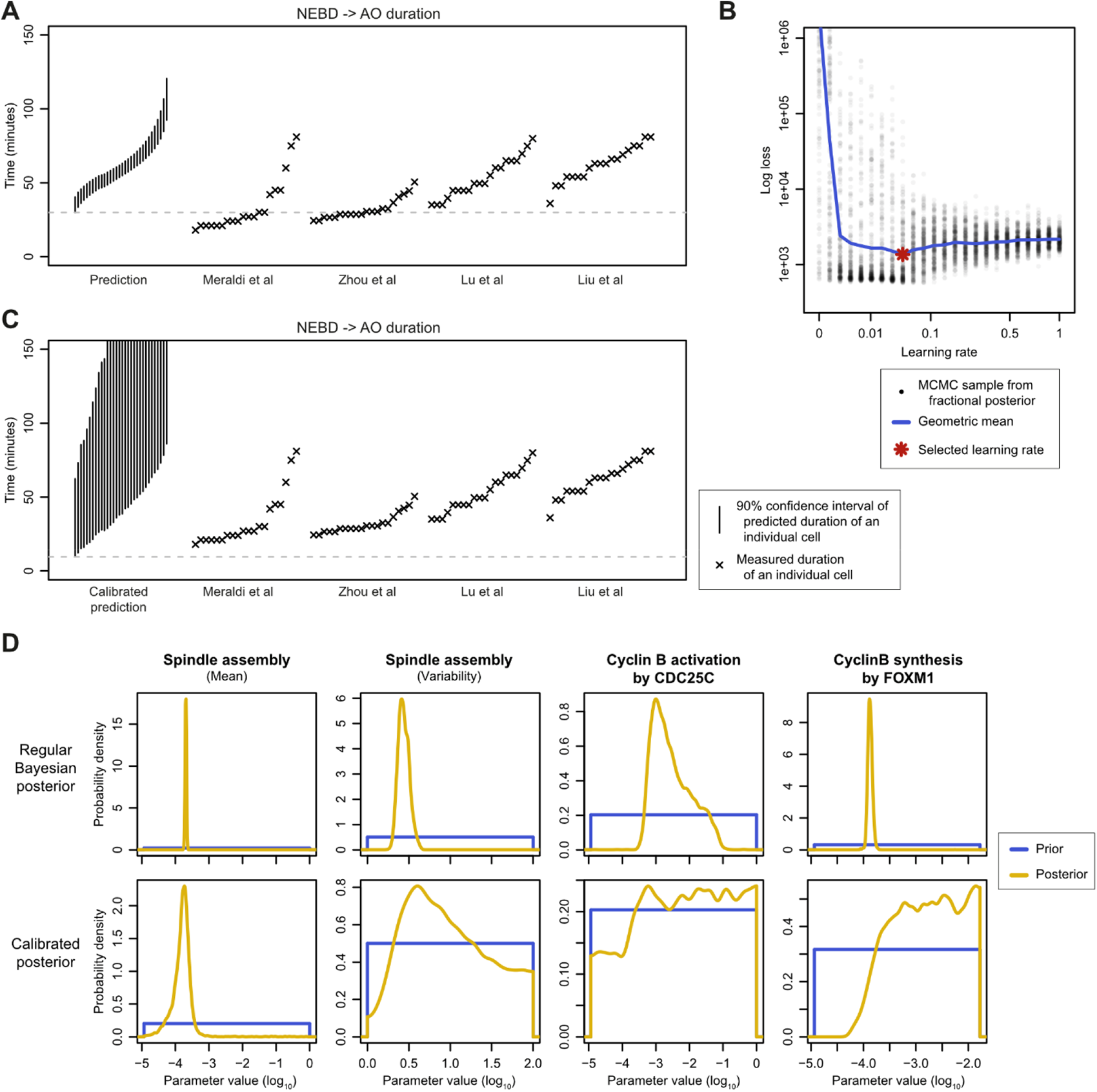
Posterior calibration to obtain robust uncertainty estimates. **(A)** Predictions of the duration from nuclear envelope breakdown (NEBD) to the onset of anaphase (AO), based on the fit to the measurements of Gavet & Pines (Figure 2), compared to four external sets of measurements of these durations. Vertical bars indicate the 90% confidence interval for every individual cell. **(B)** Posterior calibration curve used to select the calibrated posterior distribution; that is, the fractional posterior distribution that gives the minimum mean log loss in the external duration measurements. Each grey dot represents one sample from the Markov chain targeting the fractional posterior distribution at the denoted learning rate, and corresponds to the summed log loss of the four duration measurements; based on the fit to the measurements of Gavet & Pines (see Methods for details). The blue line shows the geometric mean log loss at each learning rate, and the red star marks the learning rate at which the minimum average log loss is achieved; this point gives the calibrated learning rate. **(C)** Predictions as in (A), but now with calibrated posterior confidence intervals. Some of the predicted confidence intervals extend beyond the end of the y-axis. **(D)** Comparison of the marginal posterior probability distributions of several parameters given the regular Bayesian posterior and the calibrated posterior.

This problem is consistent with previous work showing that regular Bayesian inference can lead to overconfident posterior distributions when a model is partially misspecified (Grünwald & van Ommen, 2017; Miller & Dunson, 2019). Several methods have been developed to make posterior distributions robust against misspecification (Grünwald & van Ommen, 2017; Syring & Martin, 2019; Huggins & Miller, 2023). However, these methods are far too computationally intensive when combined with lengthy simulations of populations of cells. In addition, these methods rely on the assumption that the data is independent and identically distributed. The data necessary for fitting cell cycle models are neither independent nor identically distributed: the measurements at individual time points in a time-series are not independent, and measurements from separate experiments are also not identically distributed. The measurements in separate experiments may not even describe the same physical quantities (for example, time durations and concentrations). We therefore took a more pragmatic approach and decided to use (at least) one set of measurements as inference data, and (at least) one set of orthogonal measurements as calibration data to make the posterior distributions robust against partial misspecification.

Specifically, we used a learning rate and a log-loss measure similar to what is used in SafeBayes (Grünwald & van Ommen, 2017), but rather than using forward-validation within the inference dataset as done in SafeBayes, we calculated the log loss of the entire calibration dataset. We selected the learning rate giving the minimum log loss of the calibration dataset as the learning rate for the calibrated posterior (see the Methods section for details). For the mitosis model, we used Gavet & Pines’ CDK1 sensor dataset as inference data, and the four sets of mitotic duration measurements as calibration data. Figure 3B shows the sum of the log-loss for the four calibration datasets, as a function of the learning rate. The log loss initially drops quickly as the parameter inference learns realistic parameter values, but rises again at higher learning rates when the model starts to overfit the inference data. Figure 3C shows the model prediction at the selected optimal learning rate. The uncertainty in the calibrated predictions is now much larger. With these larger confidence intervals, even the cell with the shortest duration observed in any of the external datasets still fell within the 90% confidence interval of the model predictions. Given the strong simplifications and assumptions in the model, and the limited data used for inference, it is realistic that the confidence intervals should indeed not be so small as was the case in Figure 3A, and the confidence intervals of Figure 3C are more appropriate. At the same time, the calibration did not lead to unusably wide posterior distributions. For example, the likely set of values for the average rate of spindle assembly is still constrained compared to the prior (Figure 3D). These results indicate that using a learning rate and log-loss of a set of orthogonal calibration measurements provides a method for obtaining realistic confidence intervals on the predictions in mitosis duration of individual cells.

### Calibration makes posterior distributions robust against partial misspecification

To test whether this process of posterior calibration indeed makes the posterior distributions robust against misspecification, we further investigated the resulting posteriors with a simulation study. We used the mitosis model to simulate a new dataset of CDK1 sensor measurements, using the maximum a posteriori parameter values from the fit to the Gavet & Pines data, so that we know the true parameter values for the simulated data (Figure 4A). We then fitted this simulated data with the same model that was used to generate the data, as well as with a model where we removed the positive feedback loop through CDC25C (the ‘correct’ and ‘misspecified’ models respectively, shown in Figure 4B and 4C). Without the positive feedback loop through CDC25C, the model misses an important mechanism that produces the typical switch-like activation of the cyclin B/CDK1 complex. Both the correct and the misspecified models can actually describe the data accurately (Figure 4D). However, while the misspecified model is able to describe the data, the inferred parameter values are entirely wrong (Figure 4E, second row), especially the cyclin B synthesis rate. The correct model did recover the true parameter values correctly (Figure 4E, first row). The misspecified model used the option of a very high synthesis rate of cyclin B to describe the switch-like increase of active cyclin B/CDK1 in the simulated data.

**Figure 4:**
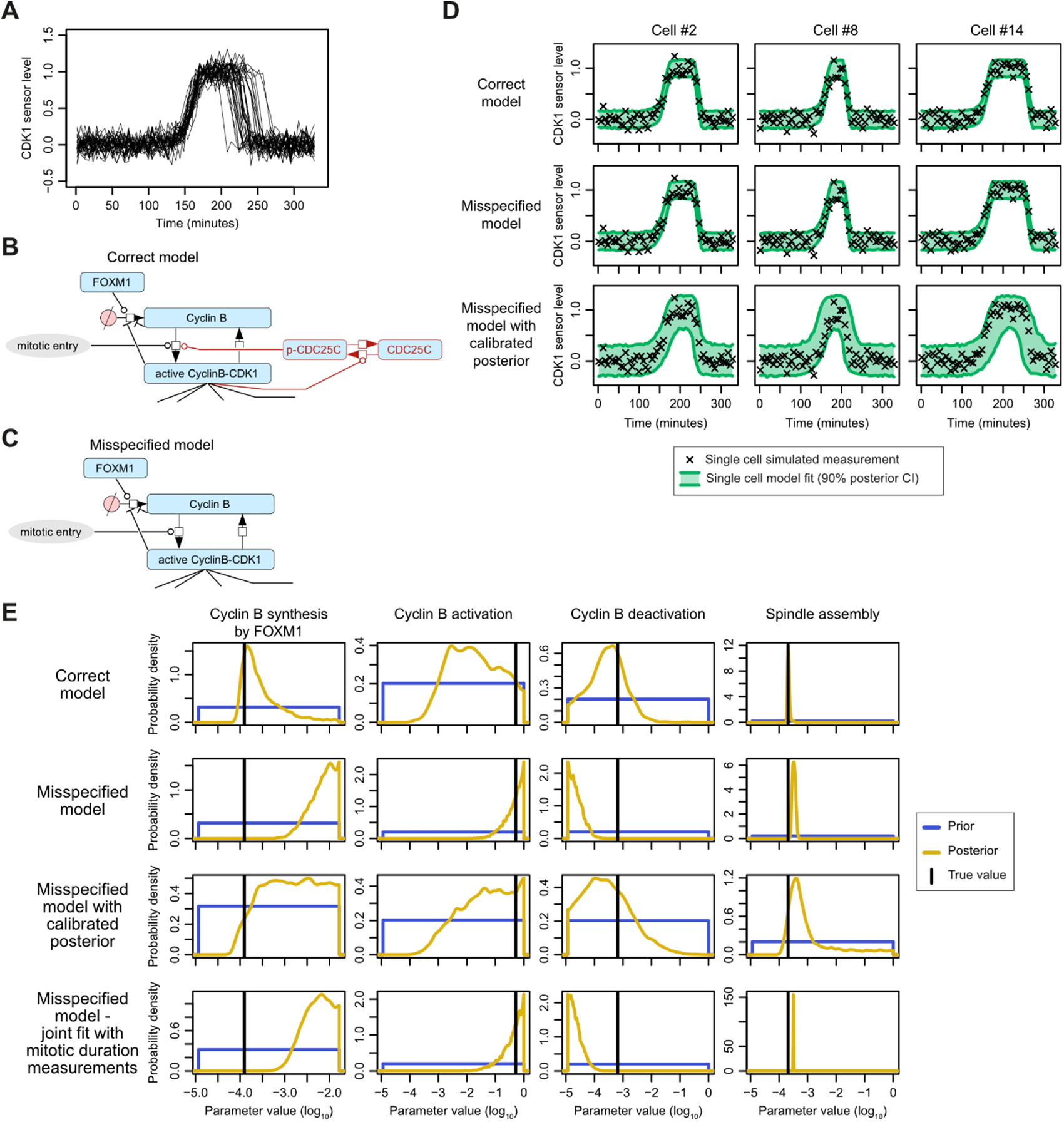
Validation of calibration process with simulated data. **(A)** A set of thirty-two CDK1 sensor traces generated from the model, given the maximum a posteriori parameter value of the fit to Gavet & Pines data, with 10% artificial noise added. **(B)** Zoom of the relevant part of the mitosis model that was used to generate the data (i.e., the correct model), the full model is shown in Figure 1A. **(C)** Zoom of the misspecified model; the positive feedback loop through CDC25C was removed. The rest of the model is identical to the correct model. The full misspecified model is shown in Supplementary Figure 2. **(D)** Model fits illustrated by three representative cells, given the correct, misspecified and calibrated misspecified model in the rows respectively; the measurements are the same for each fit. **(E)** Marginal posterior probability distributions of several parameters. The black vertical line indicates the true value that was used to generate the measurements.

We then applied the posterior calibration, which, as desired, made the uncertainty larger (Figure 4D). In the calibrated posterior distributions, the mode of the posterior distribution is still at incorrect values, but the posterior now correctly placed non-zero probability on the true values (Figure 4E, third row). That is, the parameter values that were used to generate the data are not ruled out by the calibrated posterior, whereas the non-calibrated posterior did rule out the true values. The calibrated posteriors still constrained the parameters compared to the prior distribution. Hence, the posterior calibration made the posterior distribution reliable, even though the model is partially incorrect.

The more reliable posterior distribution due to the calibration is not just an effect of adding additional data. The bottom row of Figure 4E also shows the posterior distributions that are obtained when using the external mitotic duration measurements as additional inference data rather than as calibration data. In this case, the posterior distribution is still wrong; it placed zero posterior probability on the true parameter values. Together, this shows the importance of using part of the available measurements as calibration data, rather than just using all available measurements for parameter inference.

### A simplified model of cell cycle progression can describe a variety of data

Having a methodology to simulate heterogeneous populations of cells and obtain reliable posterior distributions, we next wanted to explore if the methodology can be used to describe populations of cells going through an entire cell cycle. To this end we iteratively built a model of the cell cycle until it could describe the dynamics of various important cell cycle controls proteins. Specifically, we wanted the model to describe the dynamics of the total levels of cyclin E, cyclin A, cyclin B, geminin, p21, p27, E2F1 and E2F7, and the activity levels of CDK1 and CDK2. We started with a small model, and subsequently added regulatory mechanisms until the model could adequately describe the dynamics of these proteins and activities, in six sets of measurements obtained with a variety of measurement methodologies (Eward *et al*, 2004; Westendorp *et al*, 2012; Akopyan *et al*, 2014; Barr *et al*, 2016, 2017; Cappell *et al*, 2018; Clijsters *et al*, 2019). The resulting model is shown in Figure 5A and Supplementary Figure 3, and described in more detail in the Supplementary Materials & Methods.

**Figure 5:**
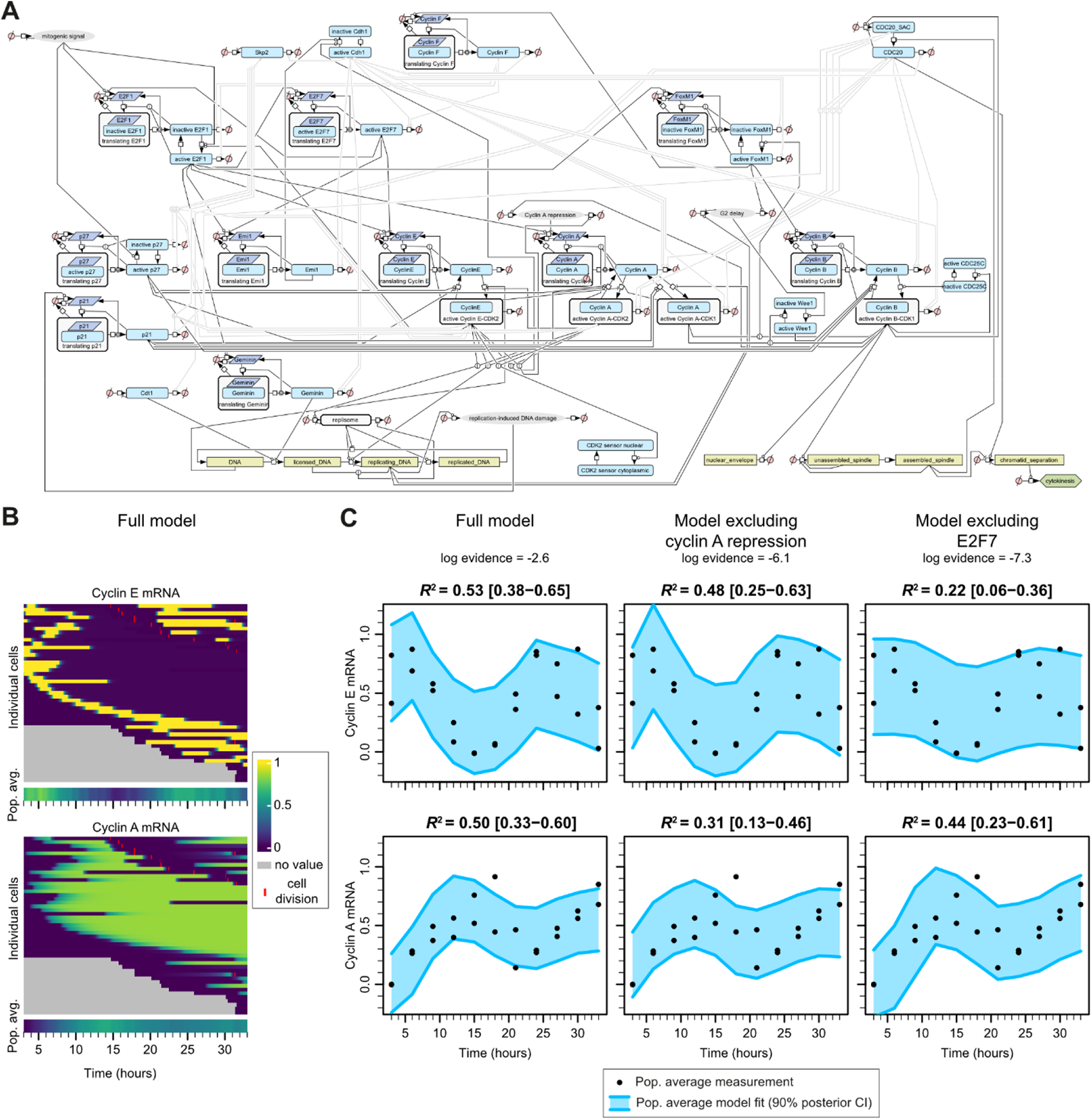
Cell cycle model and illustration of model selection with one of the sets of measurements. **(A)** Structure of the cell cycle model that is simulated in every cell in the population; a larger version is shown in Supplementary Figure 3. **(B)** Visualization of a simulated population of thirty-two initial cells and their population average, with the initial population synchronized at the start of the cell cycle. In this simulation fifteen cells divided and produced two daughter cells each. When a cell divides, one daughter takes the row of its parent and the other daughter is added at the bottom of the chart. Gray areas indicate that the cell did not exist yet. The charts show the levels of two of the dynamic variables; the mRNA levels of cyclin E and A respectively. **(C)** Measurements and posterior confidence intervals of the simulation based on the data of Ewart *et al*. (which was also synchronized at the start of the cell cycle), for three model versions. Coefficients of determination *R*^2^ were calculated as one minus the sum of squared errors given the model divided by the sum of squared errors given the data mean. Additional trajectory heatmaps and posterior predictive plots for all three models are provided in Supplementary Figure 4.

While many iterations were necessary to reach this model, we illustrate the process of model refinement with the example of cyclin E and cyclin A regulation. Cyclin E and cyclin A are both E2F target genes (Ohtani *et al*, 1995; Schulze *et al*, 1995); however cyclin A levels rise later in the cell cycle than cyclin E (Martínez-Alonso & Malumbres, 2020). To accurately describe the levels of cyclin E and cyclin A, we therefore need to include the mechanisms responsible for the delay in cyclin A expression relative to cyclin E. Specifically, it has been shown that the promotor of cyclin A contains a repressive element that needs to be relieved before cyclin A is transcribed (Schulze *et al*, 1995; Zwicker *et al*, 1995). This process was represented in the model by the Cyclin E-CDK2 complex removing the repression of cyclin A. In addition to the delay in accumulation of cyclin A transcripts, cyclin A transcription is also maintained when cyclin E transcripts are already declining. This can be explained by the finding that the repressive E2F7 binds the cyclin E promoter, but not the cyclin A promoter (Di Stefano *et al*, 2003; Westendorp *et al*, 2012).

We created a model that contains both of these mechanisms, as well as two models which lacked either of the two mechanisms respectively. Simulating a population of synchronized cells with these three models, shows that the model containing both mechanisms can describe the consecutive waves of cyclin E and cyclin A transcripts, while models that miss either of these mechanisms are inadequate (Figure 5B, C). The model without the cyclin A repressive element could not adequately describe the cyclin A dynamics simultaneously with the cyclin E dynamics, and likewise, the model excluding E2F7 is not able to describe the cyclin E transcript dynamics correctly together with the cyclin A dynamics. The coefficients of determination are clearly higher for the full model, and the marginal likelihood for the full model is indeed significantly higher than the two models excluding the two mechanisms (log Bayes factors of 3.5 and 4.7 in favor of the full model, meaning that the full model is approximately 35 and 113 times more likely respectively, given these data).

Through such steps of model refinement, we came to the full model shown in Figure 5A. To simulate the variability between cells, we again assumed that the rates of several main cellular processes were most important in describing the variability between cells. These processes were, 1) the rate of mitogenic signal accumulation, 2) the rate of DNA licensing, 3) the rate of DNA replication, 4) the rate of endogenous replication stress-induced DNA damage, and 5) the rate at which cells progress through G2 (representing delay due to reasons other than endogenous replication stress-induced DNA damage). For asynchronous experiments, a sixth cell-to-cell variability, namely variability in the cell start time, was used to generate asynchronous cells. In contrast to the mitosis model described earlier, the rate of mitotic spindle assembly was assumed to be constant across cells in the current part of the analysis, as the magnitude of variability in the duration of mitosis is typically smaller than of the other cell cycle phases (Chao *et al*, 2019). Further details of the specific choices for parameter inference are described in the Supplementary Materials & Methods.

The model could describe each of six chosen datasets accurately (Figure 6 and Supplementary Figure 5). These datasets included several measurement techniques: live cell imaging of GFP-tagged proteins (Figure 6A and B), averaged immunofluorescence after live cell imaging (Figure 6C) as well as bulk measurements done through western blotting of experimentally synchronized cells (Figure 6D). In some of the measurements there is clear variability between cells; for example, in the data of Barr *et al*, 2016 (Figure 6A), the bottom-right cell has a cyclin E peak that occurred much earlier (relative to the onset of S-phase) than the peak observed in the top-right cell, and this can be recapitulated by the model. Similarly, in the data of Barr *et al*, 2017 (Figure 6B), one cell has flat levels of p21, another cell has a small increase in p21 levels which drops again after several hours, and a third cell has p21 levels that continue to rise. The model seems to be able to describe all of these variabilities between cells.

**Figure 6:**
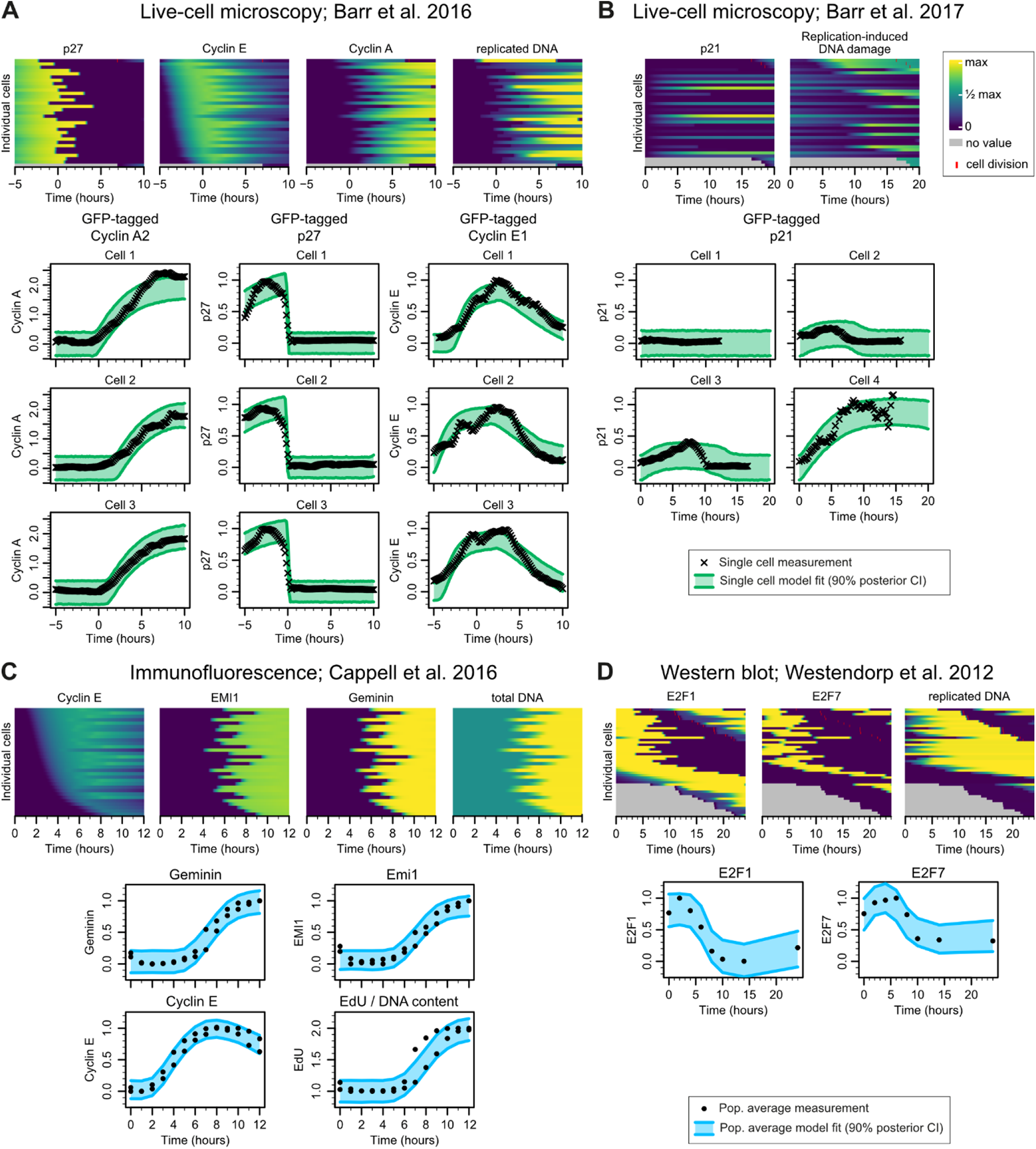
Cell cycle model simulations and fits to four additional sets of measurements. Each subfigure first shows a simulated population (given the maximum a posteriori parameter values obtained from fitting the data), followed by the 90% confidence interval of the fit to the measurements. **(A)** Measurements of endogenous GFP-tagged cyclin A, cyclin E and p27 from Barr *et al* (2016); times are relative to the start of S-phase. **(B)** Measurements of endogenous GFP-tagged p21 from Barr *et al* (2017); times are relative to the start of a cell cycle (mitosis of the parent cell). **(C)** Measurements of cyclin E, emi1, geminin and DNA content (as inferred from EdU incorporation signal) from Cappell *et al*. Immunofluorescence was obtained after live cell imaging of H2B-mTurquoise and individual cells were computationally aligned; times are relative to the start of a cell cycle (anaphase of the parent cell). **(D)** Measurements of E2F1 and E2F7 from Westendorp *et al*, through western blotting of hydroxyurea-synchronized cells; times are relative to the start of S-phase.

### The cell cycle progression model can predict levels of proteins based on CDK2 sensor measurements

One of the reasons to create these models was to use them to infer unobserved aspects of single cells, such as the levels of unmeasured proteins during live-cell imaging. To test whether the cell cycle model can make such predictions, we focused on the measurements from Gookin *et al* (Gookin *et al*, 2017). In their study, Gookin *et al* tracked MCF10A cells with a CDK2 activity sensor for 24 hours, after which the cells were fixed and the levels of one of nine proteins was measured through immunofluorescence. Six of the nine proteins that were measured were included in our model and could therefore be used as validation. To obtain single-cell predictions for these protein measurements, we used the following procedure (illustrated in Figure 7A). First, we fitted the model separately to the six inference datasets (this was described in the previous section). We then used two datasets of S-phase duration measurements (Burgess *et al*, 2012; Grant *et al*, 2018) to calibrate the posterior (see Supplementary Result 2). Next, we fixed the parameter values of the rate constants for which the six inference datasets provided information to the calibrated maximum a posteriori estimate. After this, we fitted the model to a subset of the CDK2 sensor traces of Gookin *et al*, leaving only the rates and variabilities of the main cellular processes free as inference variables, to allow the model to adapt to the MCF10A cell line. Finally, we aligned all of the CDK2 sensor traces to the simulated population, in order to predict the protein levels at the end of the trace, and compared the predictions to the observed immunofluorescence measurements. The immunofluorescence measurements were never used during inference or the construction of the model.

**Figure 7:**
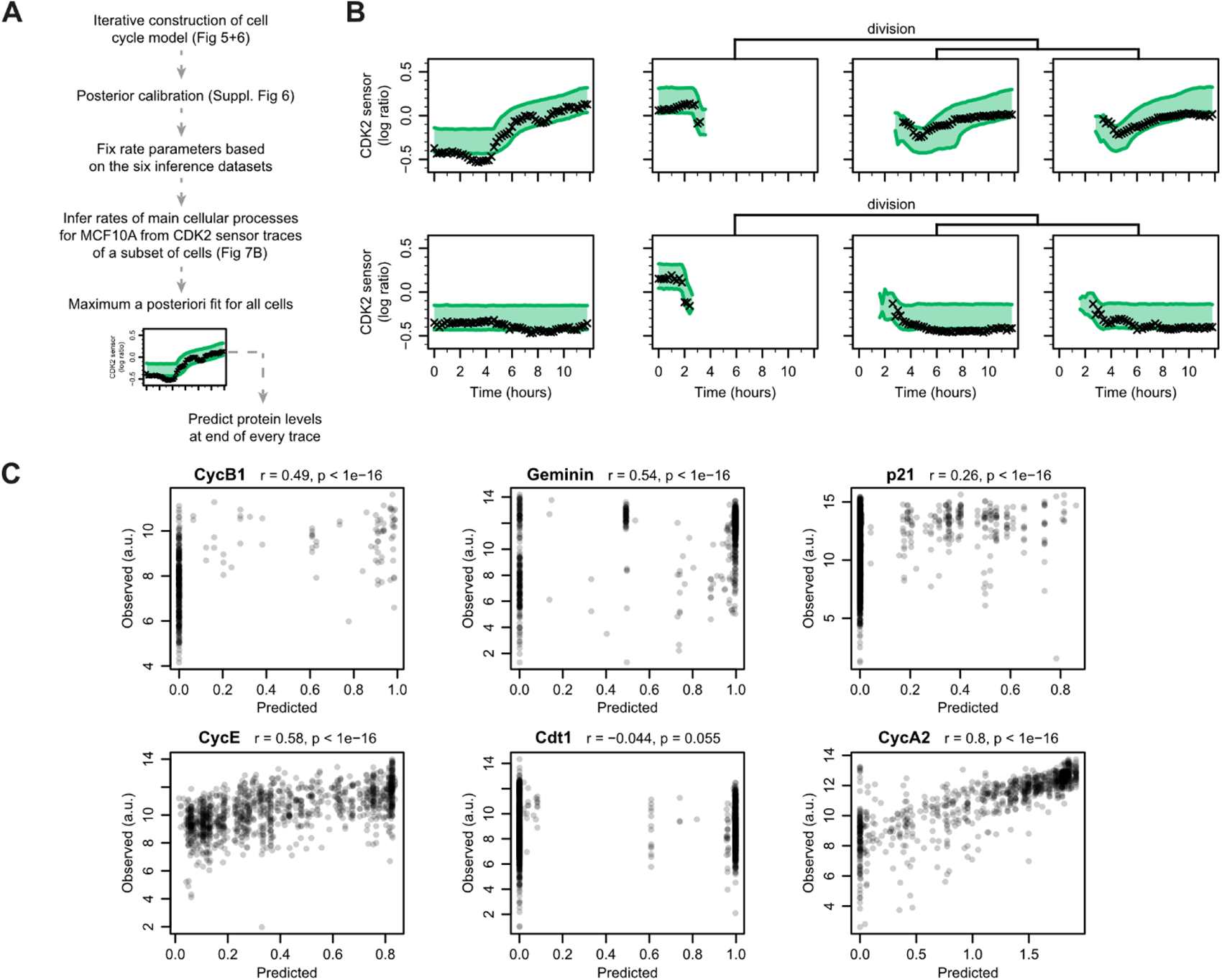
Prediction of single-cell protein levels. **(A)** Outline of the procedure to obtain single-cell protein level predictions. **(B)** Fit of the cell cycle model to CDK2 sensor measurements from the data of Gookin *et al*, allowing only the rates of the main cell cycle processes to be inferred from these CDK2 sensor traces. Four of the forty initial cells are shown, with two cells dividing into two daughter cells each. **(C)** Predicted protein levels at the end of the CDK2 trace compared to the measurements. The predicted value is the level of the protein predicted by the model for that cell, and the observed value is the level of the protein measured by immunofluorescence intensity. The r values correspond to Spearman’s rank correlation coefficient.

For inference of the main cellular process rates from the CDK2 sensor traces (that is, for the adaptation of the model to MCF10A cells), we selected 40 cells from their dataset at random (including their daughter cells if a cell divided, giving a total of 78 cell traces). We fitted these traces with a simulated population of 128 cells (growing up to 500 cells depending on cell divisions), to allow sufficient variability between the simulated cells. To reduce computation time, we simulated only the second half of the 24-hour traces. Figure 7B shows the fit to several of traces, illustrating the diversity in cell behaviors: the top left cell spends several hours in G1 and then enters S-phase (as shown by the rise in CDK2 sensor signal), the bottom left cell is arrested for the entire 12 hours, and the right cells both divided during the 12-hour window, with the two daughter cells at the top immediately progressing to S-phase with barely any time in G1, while the bottom two daughter cells both arrest at least for the 12-hour window. The simulated population can describe all these behaviors simultaneously.

To then obtain a prediction for all thousands of cells in Gookin’s set of measurements, we took the maximum a posteriori parameter value from the fit, and selected the cell from the simulation that has the best fit to each trace. The simulated protein level at the end of the trace of the selected cell was used as prediction. Figure 7C shows the comparison of the predicted protein levels plotted against the observed immunofluorescence intensities. Even though the measurements of Gookin *et al* were done in a different cell line than those used to build the model (MCF10A rather than RPE, HeLa and U2OS), five of the six proteins had a clearly significant correlation. Only one of the proteins was not predicted correctly, CDT1. It should be noted that we did not actually use any CDT1 measurements during the inference and model building, so in hindsight it is understandable that the levels of this protein could not be predicted well. In the Gookin *et al* measurements, CDT1 levels appear to drop very rapidly in newly born cells (Gookin *et al*, 2017), while the model requires some time in G1 before CDT1 is degraded. Indeed, cells may already commit to a new cell cycle before mitosis (Min *et al*, 2020), while our model always requires new mitogenic signal to be accumulated. Notwithstanding, these data show that we could accurately predict the heterogeneous levels of multiple proteins in single cells. These findings demonstrate that our computational approach can be used to infer unobserved protein levels in individual cells.

## Discussion

Understanding the molecular underpinnings of cell-to-cell variability is a major challenge. It is an important goal in -among other fields-cancer research, because cancer drug resistance often arises from subpopulations of cells within a tumor that somehow manage to continue to proliferate (Dagogo-Jack & Shaw, 2018; Marusyk *et al*, 2020). The mechanisms underlying cell decisions are complex and involve many different proteins, and computational modeling can help in understanding these complex processes.

We here presented a computational framework, scMeMo, for parameter inference of models of heterogeneous cell populations. We used this framework to construct a model describing the heterogeneous dynamics of several important cell cycle proteins during cell cycle progression. By using a parametric distribution of the variability between cells and Bayesian inference, this approach allowed transferring information between different sets of measurements, as well as making single cell predictions with reliable uncertainty estimates. A major advantage of scMeMo is that it can integrate different batches of experiments and experimental readouts, while providing biologically meaningful information about cell cycle control proteins. This is crucial, because differences between cell culture conditions, lab environments and assays can obscure true biologically relevant cell-to-cell heterogeneity (Lähnemann *et al*, 2020; Eisenstein, 2020).

The presented computational framework is an important step forward, but challenges remain. Currently we could vary relatively few parameters between cells, because otherwise prohibitively many cells need to be simulated to adequately cover the multivariate variation in parameter values. Since the computational cost scales (in the worst case) exponentially for both the number of unknown parameters as well as the number of parameters that vary between cells, there is a trade-off between these two aspects. Ways forward in this respect may be to use dedicated experiments to constrain parameters, so that available computation time can be shifted to simulating larger populations of cells rather than inferring many parameters. Alternatively, it may be possible that cell-to-cell variability can be described on a low-dimensional manifold, such that a limited number of cells can still adequately describe extensive intracellular heterogeneity.

A second limitation is that the simulations are deterministic, and stochasticity in biochemical reactions is not taken into account. Stochasticity for some reactions could still be incorporated by specifying a rate constant that varies between cells for those reactions. More extensive stochasticity however, such as bursting gene expression or stochastic intra-cellular variability in signaling pulses, cannot currently be incorporated. Other methods of parameter inference can enable inference for stochastic simulations, such as with likelihood-free methods (Frazier & Drovandi, 2021; Ward *et al*, 2022) or dynamic prior propagation (Zechner *et al*, 2014). However, the computational cost of such methods is currently prohibitive for extensive models of signaling networks like the cell cycle model we investigated here.

Despite these limitations, the advantage of the assumptions we made is that the populations of cells can be simulated sufficiently fast to enable elaborate statistical inference. Specifically, the framework allowed for multivariate parameter inference, with multiple datasets, such that models could be constrained with several complementary types of measurements. It furthermore allowed for posterior calibration to make uncertainty estimates robust against partial model misspecification. As a result, the presented modeling approach can be used to integrate bulk and single-cell measurements, and to make single-cell inferences even when cells in a population differ from each other.

## Materials and Methods

### Terminology

For reference, Table 1 specifies the symbols and meanings for specific terms used in this manuscript. Throughout the formulas, the indices *i* enumerate cells, *j* enumerates dynamic variables, *k* enumerates model parameters and *l* enumerates inference variables. In the equations, bold letters indicate vectors and italic letters indicate scalars.

**Table 1:**
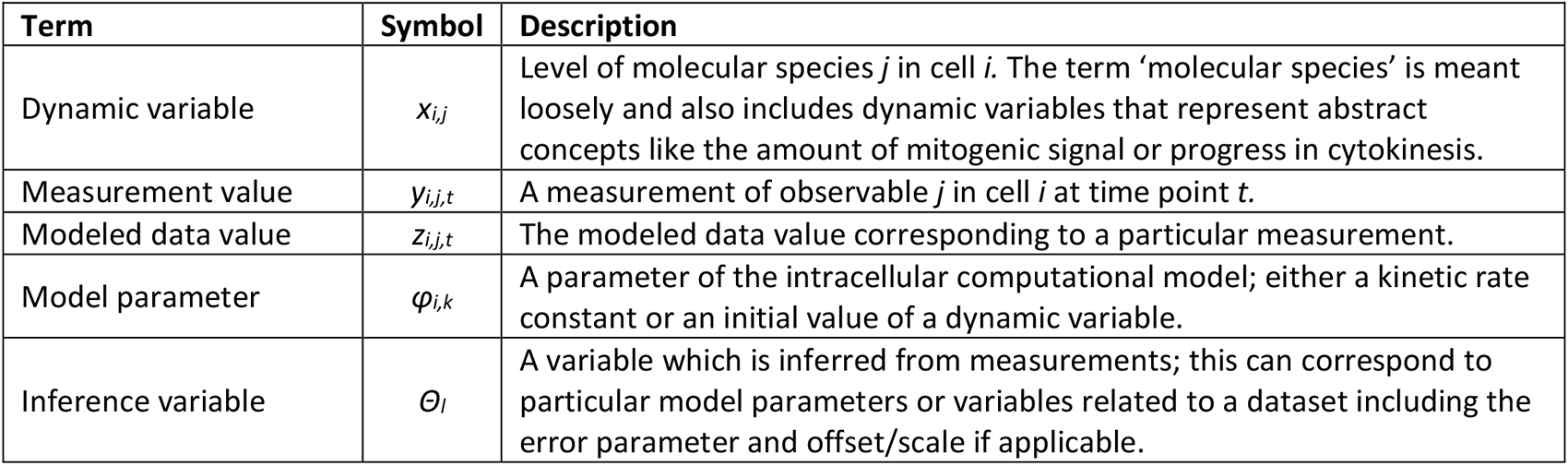
Terms and symbols used in the text and equations.

### Cell populations

The population of cells is simulated as a set of ordinary differential equation systems. Each cell in the population uses the same model structure, but has a separate instance of the ODE system. The dynamics of the population of cells is described by the differential equation

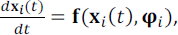

where **x***_i_* is the vector of molecular species levels in cell *i*, and φ*_i_* is the vector of model parameters in cell *i*, which includes both kinetic rate constants and initial values. There are two scenarios for determining the initial conditions of each molecular specie in a cell. At the start of the simulation, the initial conditions are specified as part of the model, or can be included as inference variable. Alternatively, when new daughter cells are generated through cytokinesis, the initial conditions are copied over from the mother cell, with some specific exceptions (for instance, the amount of DNA is divided in half for each daughter cell; see Supplementary Materials & Methods for details). The function **f** is determined by the rate laws of the reactions in the model; these rate laws are specified in the SBML file describing the model. For the two models described in this manuscript, we used a combination of mass action kinetics, Michaelis-Menten and hill kinetics, as described in more detail in the Supplementary Materials & Methods.

A model can contain a dynamic variable for cytokinesis. When this is the case, this variable will be used to control the generation of new daughter cells. When the cytokinesis variable reaches a value of 1 during the simulation, the parent cell simulation is stopped, and two new daughter cells are generated and added to the set of cells. A dynamic variable for apoptosis can also be included in the model, which will stop the simulation of that cell without generating new children cells, when it reaches a value of 1.

The ODE systems were integrated using an optimized version of the CVODE algorithm of the SUNDIALS package (Hindmarsh *et al*, 2005; Gardner *et al*, 2022). The model structures and rate equations were specified using CellDesigner (Funahashi *et al*, 2003) and stored in SBML format, the data sets are stored in NetCDF files, and the priors and likelihoods are specified in XML description files. All code for simulation of scMeMo models, as well as the calculation of the likelihoods described below, are included in a module “cellpop” included in version 3 of BCM (Thijssen *et al*, 2016), available at https://github.com/NKI-CCB/bcm3.

### Variability between cells

To keep the computational burden manageable, we set most kinetic rate parameters and initial values to be the same for all cells; that is, most elements of φ*_i_* are the same for all cells. Only some of the model parameters are allowed to vary between cells. The decision of which parameters are allowed to vary depends on the research question the model is used for. For the mitosis model and cell cycle model these decisions are described in the results section.

When a parameter varies between cells, we assume that the rates are normally distributed on a log_2_ scale:

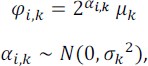

where *μ_k_* indicates the mean parameter value and *σ_k_* indicates the extent of variability for parameter *k*. *μ_k_* and *σ_k_* are typically included as unknowns in the inference variables **θ**. If so, *μ_k_* is given the same prior as when it is a parameter that does not vary between cells, and *σ_k_* is given a uniform prior between 0 and 2, so that the resulting variabilities can be anything between no variability at all and an approximately 100-fold difference between the 5^th^ and 95^th^ percentile across cells. To keep the simulations deterministic, the cell-specific values for these varying parameters are drawn from the normal distribution through a Sobol sequence with dimension equal to the number of parameters that vary between cells.

### Likelihood

Since we want to use the model to make predictions between distinct sets of measurements, we need a way to set the expected prediction error of the unseen set of measurements. However, the types of measurements can be entirely different physical quantities (for example, we wanted to predict the duration of a cell cycle phase, based on measurements of protein levels, meaning we have to go from concentration measurements to time measurements). To accommodate this, we do not use a specific standard deviation of the error for each dataset, but describe the error relative to the observed dispersion of the measurements. Specifically, the standard deviation of the error is specified as a fraction of the interquartile range of the measurements, and the following likelihood function is used:

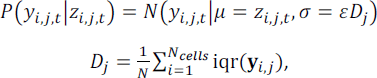

where *y_i,j,t_* is the observed value for observable *j* in cell *i* at timepoint *t*, z*_i,j,t_* is the modeled data value for observable *j* in cell *i* at timepoint *t*, *ε* is the fractional error term and **y***_i,j_* indicates the vector of the entire observed time course for observable *j* in cell *i*. As described in the equation, when the measurements are cell-specific, we took the mean of the interquartile ranges of the individual cells. When the measurements come from bulk cells, we took the single interquartile range of the bulk time course measurement (if there were no replicate measurements), or the average interquartile ranges over biological replicates (if there were replicates). When the set of measurements is cross-sectional rather than longitudinal, we take the interquartile range of the cross-section. When fitting to an average over single-cell measurements, the standard deviation of the error is divided by the square root of the number of cells being averaged over. An ε with a value of 1 indicates an error equal to the dispersion of the data, meaning that the model is not able to describe the data at all, whereas an ε approaching 0 indicates the model can describe the data very accurately. Using this setup, the model uncertainty can be transferred between distinct sets of data; when a model can describe a particular dataset reasonably well, we would *a priori* expect the uncertainty in other datasets to be on a similar level, whereas when a model cannot describe a dataset well, we would not expect it to predict other datasets well either. We included this fractional error parameter as inference variable, on a log_10_ scale with a uniform prior with lower and upper bounds of -2 and 0, corresponding to an error between 1% and 100% relative to the interquartile range.

Since all measurements of molecular species that were used provided relative values, we normalized all measurements between 0 and 1, to match them with the dynamic variables of the kinetic model that were also constrained between 0 and 1. When a measurement directly corresponds to a dynamic variable, the modeled data value ***z*** is set directly equal to the value of the corresponding dynamic variable ***y*** at that time point. In some cases a measurement does not directly correspond to the level of one molecular species in the model; for example, a measurement of the total amount of cyclin E would include both of the dynamic variables inactive Cyclin E and active Cyclin E in complex with CDK2. In these cases the modeled data values ***z*** are set to the sum of relevant dynamic variables. Likewise a biosensor measurement may be a log ratio between two dynamic variables, in which case the modeled data value is also calculated as a log ratio. For measurements based on antibodies, including western blots and immunofluorescence, we further included an offset and scale parameter, which were included as inference variables, to allow for signal from background staining.

Since we simulated populations of cells, we had to match individual simulated cells to individual observed cells. To do this, we calculated the likelihood for all possible pairs of simulated and observed cells, and used the Hungarian matching algorithm to select the optimal matching. When the time course data included information on cell divisions and tracked the daughter cells belonging to a specific mother cell, we matched the entire hierarchy of mother/daughter cells between the simulations and observations. This can lead to situations where there is no simulated data for a particular measurement, when a simulated cell does not divide while the observed cell does. We penalized such a matching based on the amount of time during which matchings cannot be made because of a lack of cell division, with a normal error model with a standard deviation that is either fixed or included as inference variables. For bulk measurements, we calculated the average of all cells in the simulated population at a specific time point. If the bulk or averaged measurements contained selection (for instance, the bulk measurement may include only cells that went through mitosis); then we did the same selection in the simulation).

### Parameter inference

We used Bayesian inference to estimate the parameter values and their uncertainty from the measurements. When a parameter or parameter variability was set to be an inference variables, the prior for it was set to a weakly informative uniform distribution, with upper and lower bounds based on biochemically plausible limits (specified in Supplementary Table 1 and 2 for the mitosis and full cell cycle model respectively).

The posterior distributions were estimated using parallel-tempered Markov chain Monte Carlo (MCMC) sampling (Earl & Deem, 2005). The parallel-tempering benefits the convergence to the target MCMC chain by providing a way to move between local optima, but also directly gives a range of fractional posterior distributions that are used for the posterior calibration process described in the next section.

In addition to the parallel-tempering, an efficient proposal distribution for mutation moves was needed, as the posterior distributions that arose with the present model and data often contained ridges and modes that differed in shape. We therefore implemented a novel locally adaptive proposal distribution based on shrunken Gaussian mixtures to efficiently sample from the posterior distributions (described in more detail in the Supplementary Materials & Methods). This sampling scheme was added to version 3 of the BCM sampling software so that it can be used for other inference problems as well (available at https://github.com/NKI-CCB/bcm3). Convergence was assessed by visual inspection of the posterior sample traces, and the number of samples was increased as necessary until autocorrelation patterns in the sample traces were minimal.

### Posterior calibration

The posterior calibration was inspired by the SafeBayes procedure (Grünwald & van Ommen, 2017). However, SafeBayes assumes that the data under consideration are independent and identically distributed. This means SafeBayes cannot be directly applied here, as longitudinal data are not independent, and the separate datasets are not identically distributed. We therefore cannot build on the theoretical guarantees of SafeBayes. Instead, we just assume that the log-loss of an orthogonal dataset is a good quantity to optimize in order to calibrate the posterior. Instead of the point-wise forward-validation, we calculated the log-loss of all external validation datasets based on the fractional posteriors of the inference dataset. These fractional posteriors were directly available from the parallel tempering scheme used during the sampling procedure. We then selected the learning rate that gave the lowest mean log-loss of the validation datasets, and used the fractional posterior corresponding to that learning rate as the calibrated posterior.

The fractional posterior distributions are given by

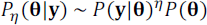

where *η* is the learning rate (i.e., the ‘fraction’ in the term ‘fractional posterior’), **y** is the set of measurements and **θ** are the parameters to be inferred. We evaluated the log loss of the external validation sets and selected the optimal learning rate by:

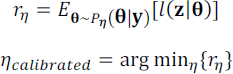

where **z** are the external validation measurements, *l* is the log loss, where the log loss is set to the negative log likelihood of the validation measurements, and *r_η_* is the expected log-loss at learning rate *η*, calculated as the average log loss given the posterior samples obtained from the inference dataset at that learning rate. The present models are highly nonlinear, and the distribution of log likelihood values can be highly skewed; we therefore used the geometric mean to estimate the expected log-loss.

### External data

Where possible, data was obtained from supplementary information of the corresponding publications. When the raw data was not provided, we derived the data from the figures as follows. When the figures were included in the publication as a vector graphic, we exported the figure to SVG format, imported them to R and regressed the data points against the tick marks to obtain the measurement values.

When the figures were in bitmap format, we traced the points or lines and the tick marks manually, exported the traces to SVG format and used the same regression procedure.

### Nuclear envelope breakdown to anaphase onset duration measurement

We measured the duration of nuclear envelope breakdown until the onset of anaphase in HeLa cells. These measurements were part of another investigation into atypical E2F transcription factors (Liu et al, manuscript in revision), but are included here as well. HeLa Cells were transduced with the lentiviral expression vector pLenti-H2B-iRFP720 (a gift from Carlos Carmona-Fontaine (Addgene plasmid # 128961; RRID:Addgene_128961) to express fluorescently labeled H2B proteins according to methodology described previously (Segeren *et al*, 2022). Stable expression was achieved by 3 days of Zeocin selection. Twenty-five thousand cells were plated in 4-well CELLview culture dishes (627975, Greiner), and transfected with scrambled siRNA after one day. Twenty-four hours after siRNA transfection, cells were treated with 0.2 mM thymidine for 16 hours. Cells were subjected to imaging after 8 hours of thymidine release. Images were acquired every 3 minutes for 6 hours on a Nikon A1R-STORM microscope using a 10x objective in a humidified chamber at 37℃, 5% CO2. Cell tracking was performed manually with NIS Elements software.

## Data Availability

The scMeMo simulation and Bayesian inference code is available in the BCM software package version 3 available at https://github.com/NKI-CCB/bcm3. All other code used in the analysis in this paper is provided at https://github.com/NKI-CCB/scMeMo-paper.

## Acknowledgements

We thank Richard Wubbolts (Center for Cellular Imaging, Utrecht University) for support with live cell imaging. We thank Alain de Bruin for constructive feedback and discussions. This work made use of the Dutch national e-infrastructure with the support of the SURF Cooperative using grant no. EINF-2116 and EINF-5932. This research was supported in part by the Dutch Cancer Society (project grant 11941-2018-II), and by institutional grants of the Dutch Cancer Society and Dutch Ministry of Health, Welfare and Sport to the Netherlands Cancer Institute.

## Author contributions

**Bram Thijssen:** Conceptualization, Data curation, Funding Acquisition, Formal Analysis, Investigation, Methodology, Software, Validation, Visualization, Writing – original draft, Writing – review & editing. **Hendrika A Segeren:** Conceptualization, Methodology, Writing – review & editing. **Qingwu Liu:** Investigation. **Lodewyk FA Wessels:** Conceptualization, Funding Acquisition, Methodology, Supervision, Writing – review & editing. **Bart Westendorp:** Conceptualization, Funding Acquisition, Methodology, Supervision, Writing – review & editing.

## Disclosure and competing interests statement

The authors declare that they have no conflict of interest.

## Notes

https://github.com/NKI-CCB/scMeMo-paper

### Competing Interest Statement

The authors have declared no competing interest.

## References

Akopyan K, Silva Cascales H, Hukasova E, Saurin AT, Müllers E, Jaiswal H, Hollman DAA, Kops GJPL, Medema RH & Lindqvist A (2014) Assessing Kinetics from Fixed Cells Reveals Activation of the Mitotic Entry Network at the S/G2 Transition. Molecular Cell 53: 843–853

Bajar BT, Lam AJ, Badiee RK, Oh Y-H, Chu J, Zhou XX, Kim N, Kim BB, Chung M, Yablonovitch AL, et al. (2016) Fluorescent indicators for simultaneous reporting of all four cell cycle phases. Nature Methods 13: 993–996

Bandura DR, Baranov VI, Ornatsky OI, Antonov A, Kinach R, Lou X, Pavlov S, Vorobiev S, Dick JE & Tanner SD (2009) Mass Cytometry: Technique for Real Time Single Cell Multitarget Immunoassay Based on Inductively Coupled Plasma Time-of-Flight Mass Spectrometry. Anal Chem 81: 6813–6822

Barr AR, Cooper S, Heldt FS, Butera F, Stoy H, Mansfeld J, Novák B & Bakal C (2017) DNA damage during S-phase mediates the proliferation-quiescence decision in the subsequent G1 via p21 expression. Nat Commun 8: 14728

Barr AR, Heldt FS, Zhang T, Bakal C & Novák B (2016) A Dynamical Framework for the All-or-None G1/S Transition. Cell Systems 2: 27–37

Bhuyan BK, Scheldt LG & Fraser TJ (1972) Cell Cycle Phase Specificity of Antitumor Agentsl. Cancer Research 32: 398–407

Browning AP, Ansari N, Drovandi C, Johnston APR, Simpson MJ & Jenner AL (2022a) Identifying cell-to-cell variability in internalization using flow cytometry. J R Soc Interface 19: 20220019

Browning AP, Drovandi C, Turner IW, Jenner AL & Simpson MJ (2022b) Efficient inference and identifiability analysis for differential equation models with random parameters. PLoS Comput Biol 18: e1010734

Burgess A, Lorca T & Castro A (2012) Quantitative Live Imaging of Endogenous DNA Replication in Mammalian Cells. PLoS ONE 7: e45726

Cappell SD, Mark KG, Garbett D, Pack LR, Rape M & Meyer T (2018) EMI1 switches from being a substrate to an inhibitor of APC/CCDH1 to start the cell cycle. Nature 558: 313–317

Chao HX, Fakhreddin RI, Shimerov HK, Kedziora KM, Kumar RJ, Perez J, Limas JC, Grant GD, Cook JG, Gupta GP, et al. (2019) Evidence that the human cell cycle is a series of uncoupled, memoryless phases. Mol Syst Biol 15

Clijsters L, Hoencamp C, Calis JJA, Marzio A, Handgraaf SM, Cuitino MC, Rosenberg BR, Leone G & Pagano M (2019) Cyclin F Controls Cell-Cycle Transcriptional Outputs by Directing the Degradation of the Three Activator E2Fs. Molecular Cell 74: 1264–1277.e7

Dagogo-Jack I & Shaw AT (2018) Tumour heterogeneity and resistance to cancer therapies. Nat Rev Clin Oncol 15: 81–94

Dharmarajan L, Kaltenbach H-M, Rudolf F & Stelling J (2019) A Simple and Flexible Computational Framework for Inferring Sources of Heterogeneity from Single-Cell Dynamics. Cell Systems 8: 15–26.e11

Di Stefano L, Jensen MR & Helin K (2003) E2F7, a novel E2F featuring DP-independent repression of a subset of E2F-regulated genes. The EMBO Journal 22: 6289–6298

Dixit PD, Lyashenko E, Niepel M & Vitkup D (2020) Maximum Entropy Framework for Predictive Inference of Cell Population Heterogeneity and Responses in Signaling Networks. Cell Systems 10: 204–212.e8

Earl DJ & Deem MW (2005) Parallel tempering: Theory, applications, and new perspectives. Phys Chem Chem Phys 7: 3910

van Eijl RAPM, van Buggenum JAGL, Tanis SEJ, Hendriks J & Mulder KW (2018) Single-Cell ID-seq Reveals Dynamic BMP Pathway Activation Upstream of the MAF/MAFB-Program in Epidermal Differentiation. iScience 9: 412–422

Eisenstein M (2020) Single-cell RNA-seq analysis software providers scramble to offer solutions. Nat Biotechnol 38: 254–257

Eward KL, Ert MNV, Thornton M & Helmstetter CE (2004) Cyclin mRNA Stability Does Not Vary During the Cell Cycle. Cell Cycle 3: 1055–1059

Frazier DT & Drovandi C (2021) Robust Approximate Bayesian Inference With Synthetic Likelihood. Journal of Computational and Graphical Statistics 30: 958–976

Funahashi A, Morohashi M, Kitano H & Tanimura N (2003) CellDesigner: a process diagram editor for gene-regulatory and biochemical networks. BIOSILICO 1: 159–162

Gardner DJ, Reynolds DR, Woodward CS & Balos CJ (2022) Enabling New Flexibility in the SUNDIALS Suite of Nonlinear and Differential/Algebraic Equation Solvers. ACM Trans Math Softw 48: 1–24

Gavet O & Pines J (2010) Progressive Activation of CyclinB1-Cdk1 Coordinates Entry to Mitosis. Developmental Cell 18: 533–543

Gookin S, Min M, Phadke H, Chung M, Moser J, Miller I, Carter D & Spencer SL (2017) A map of protein dynamics during cell-cycle progression and cell-cycle exit. PLoS Biol 15: e2003268

Grant GD, Kedziora KM, Limas JC, Cook JG & Purvis JE (2018) Accurate delineation of cell cycle phase transitions in living cells with PIP-FUCCI. Cell Cycle 17: 2496–2516

Greenberg DS, Nonnenmacher M & Macke JH (2019) Automatic Posterior Transformation for Likelihood-free Inference. In Proceedings of the 36th International Conference on Machine Learning pp 2404–2414.

Grünwald P & van Ommen T (2017) Inconsistency of Bayesian Inference for Misspecified Linear Models, and a Proposal for Repairing It. Bayesian Anal 12

Gut G, Herrmann MD & Pelkmans L (2018) Multiplexed protein maps link subcellular organization to cellular states. Science 361: eaar7042

Hindmarsh AC, Brown PN, Grant KE, Lee SL, Serban R, Shumaker DE & Woodward CS (2005) SUNDIALS: Suite of nonlinear and differential/algebraic equation solvers. ACM Trans Math Softw 31: 363– 396

Hsu C-H, Altschuler SJ & Wu LF (2019) Patterns of Early p21 Dynamics Determine Proliferation-Senescence Cell Fate after Chemotherapy. Cell 178: 361–373.e12

Huggins JH & Miller JW (2023) Reproducible Model Selection Using Bagged Posteriors. Bayesian Anal 18

Kolodziejczyk AA, Kim JK, Svensson V, Marioni JC & Teichmann SA (2015) The Technology and Biology of Single-Cell RNA Sequencing. Molecular Cell 58: 610–620

Lähnemann D, Köster J, Szczurek E, McCarthy DJ, Hicks SC, Robinson MD, Vallejos CA, Campbell KR, Beerenwinkel N, Mahfouz A, et al. (2020) Eleven grand challenges in single-cell data science. Genome Biol 21: 31

Lambert B, Gavaghan DJ & Tavener SJ (2021) A Monte Carlo method to estimate cell population heterogeneity from cell snapshot data. Journal of Theoretical Biology 511: 110541

Laoukili J, Stahl M & Medema RH (2007) FoxM1: At the crossroads of ageing and cancer. Biochimica et Biophysica Acta (BBA) – Reviews on Cancer 1775: 92–102

Letort G, Montagud A, Stoll G, Heiland R, Barillot E, Macklin P, Zinovyev A & Calzone L (2019) PhysiBoSS: a multi-scale agent-based modelling framework integrating physical dimension and cell signalling. Bioinformatics 35: 1188–1196

Liao G-B, Li X-Z, Zeng S, Liu C, Yang S-M, Yang L, Hu C-J & Bai J-Y (2018) Regulation of the master regulator FOXM1 in cancer. Cell Commun Signal 16: 57

Loos C, Moeller K, Fröhlich F, Hucho T & Hasenauer J (2018) A Hierarchical, Data-Driven Approach to Modeling Single-Cell Populations Predicts Latent Causes of Cell-To-Cell Variability. Cell Systems 6: 593–603.e13

Lu L, Hu S, Wei R, Qiu X, Lu K, Fu Y, Li H, Xing G, Li D, Peng R, et al. (2013) The HECT Type Ubiquitin Ligase NEDL2 Is Degraded by Anaphase-promoting Complex/Cyclosome (APC/C)-Cdh1, and Its Tight Regulation Maintains the Metaphase to Anaphase Transition. Journal of Biological Chemistry 288: 35637–35650

Martínez-Alonso D & Malumbres M (2020) Mammalian cell cycle cyclins. Seminars in Cell & Developmental Biology 107: 28–35

Marusyk A, Janiszewska M & Polyak K (2020) Intratumor Heterogeneity: The Rosetta Stone of Therapy Resistance. Cancer Cell 37: 471–484

Meraldi P, Draviam VM & Sorger PK (2004) Timing and Checkpoints in the Regulation of Mitotic Progression. Developmental Cell 7: 45–60

Miller JW & Dunson DB (2019) Robust Bayesian Inference via Coarsening. Journal of the American Statistical Association 114: 1113–1125

Millour J, de Olano N, Horimoto Y, Monteiro LJ, Langer JK, Aligue R, Hajji N & Lam EW-F (2011) ATM and p53 Regulate FOXM1 Expression via E2F in Breast Cancer Epirubicin Treatment and Resistance. Molecular Cancer Therapeutics 10: 1046–1058

Min M, Rong Y, Tian C & Spencer SL (2020) Temporal integration of mitogen history in mother cells controls proliferation of daughter cells. Science 368: 1261–1265

Ohtani K, DeGregori J & Nevins JR (1995) Regulation of the cyclin E gene by transcription factor E2F1. Proc Natl Acad Sci USA 92: 12146–12150

Papamakarios G, Sterratt DC & Murray I (2019) Sequential Neural Likelihood: Fast Likelihood-free Inference with Autoregressive Flows. In Proceedings of the Twenty-Second International Conference on Artificial Intelligence and Statistics pp 837–848.

Schulze A, Zerfass K, Spitkovsky D, Middendorp S, Bergès J, Helin K, Jansen-Dürr P & Henglein B (1995) Cell cycle regulation of the cyclin A gene promoter is mediated by a variant E2F site. Proc Natl Acad Sci USA 92: 11264–11268

Segeren HA, van Liere EA, Riemers FM, de Bruin A & Westendorp B (2022) Oncogenic RAS sensitizes cells to drug-induced replication stress via transcriptional silencing of P53. Oncogene

Segeren HA, van Rijnberk LM, Moreno E, Riemers FM, van Liere EA, Yuan R, Wubbolts R, de Bruin A & Westendorp B (2020) Excessive E2F Transcription in Single Cancer Cells Precludes Transient Cell-Cycle Exit after DNA Damage. Cell Reports 33: 108449

Shaltiel IA, Krenning L, Bruinsma W & Medema RH (2015) The same, only different – DNA damage checkpoints and their reversal throughout the cell cycle. Journal of Cell Science: jcs.163766

Spencer SL, Gaudet S, Albeck JG, Burke JM & Sorger PK (2009) Non-genetic origins of cell-to-cell variability in TRAIL-induced apoptosis. Nature 459: 428–432

Syring N & Martin R (2019) Calibrating general posterior credible regions. Biometrika 106: 479–486

Thijssen B, Dijkstra TMH, Heskes T & Wessels LFA (2016) BCM: toolkit for Bayesian analysis of Computational Models using samplers. BMC Syst Biol 10: 100

Ward D, Cannon P, Beaumont M, Fasiolo M & Schmon S (2022) Robust Neural Posterior Estimation and Statistical Model Criticism. In Advances in Neural Information Processing Systems, Koyejo S Mohamed S Agarwal A Belgrave D Cho K & Oh A (eds) pp 33845–33859. Curran Associates, Inc.

Westendorp B, Mokry M, Groot Koerkamp MJA, Holstege FCP, Cuppen E & de Bruin A (2012) E2F7 represses a network of oscillating cell cycle genes to control S-phase progression. Nucleic Acids Research 40: 3511–3523

Zechner C, Unger M, Pelet S, Peter M & Koeppl H (2014) Scalable inference of heterogeneous reaction kinetics from pooled single-cell recordings. Nat Methods 11: 197–202

Zhou L, Liang C, Chen Q, Zhang Z, Zhang B, Yan H, Qi F, Zhang M, Yi Q, Guan Y, et al. (2017) The N-Terminal Non-Kinase-Domain-Mediated Binding of Haspin to Pds5B Protects Centromeric Cohesion in Mitosis. Current Biology 27: 992–1004

Zwicker J, Lucibello FC, Wolfraim LA, Gross C, Truss M, Engeland K & Müller R (1995) Cell cycle regulation of the cyclin A, cdc25C and cdc2 genes is based on a common mechanism of transcriptional repression. The EMBO Journal 14: 4514–4522

