## supplementary-information for "Inferring single-cell protein levels and cell cycle behavior in heterogeneous cell populations"

#### Model description

A short overview of both the mitosis and cell cycle model is provided first, followed by a more detailed description of specific parts of the models.

##### Mitosis model overview

The mitosis model is meant as a simple test case with a minimal amount of mechanistic detail necessary to describe several main events of mitosis. Specifically, we wanted the model to describe the point at which the nuclear envelope is broken down, the assembly of the mitotic spindle, and the separation of the sister chromatids. We assumed that nuclear envelope breakdown and synthesis of the spindle components are both triggered when active cyclin B-CDK1 complexes reach the arbitrary level of 0.5. The spindle is then assembled with quasi-zero order kinetics. When the spindle is fully assembled, the spindle assembly checkpoint is relieved, modeled by the conversion of CDC20_with_SAC into free CDC20.

In the model, cyclin B is synthesized under the control of FOXM1. A mitotic entry trigger needs to accumulate to the arbitrary value of 0.5 before cyclin B starts to form an active complex with CDK1. This trigger could represent the removal of Wee1/Myt1 activity, among others. One positive feedback is included, through CDC25C, to provide a switch-like entry into mitosis. Finally, we incorporated an interaction that CDC20 degrades FOXM1 and stops its production. This does not represent known biology, but is an artificial interaction to prevent re-entry into mitosis. This is necessary since the cells in the model don't actually divide; and something has to stop FOXM1 becoming active and triggering an immediate mitosis again.

The CDK1 sensor is included as three dynamic variables, one variable each for the inactive and active variant, and a variable representing dilution. The inactive variant is converted into the active variant by the active cyclin B-CDK1 complex. Gavet & Pines observed a decrease in sensor signal during and after mitosis, which was attributed to differential bleaching of the FRET moieties and rounding of the cells. This decrease in sensor signal was incorporated into the model as a dilution value which decreased the signal after the nuclear envelope was broken down.

This model is a major simplification of the complex regulation of mitotic processes. The goal of the model is not to describe mitosis in large detail but rather to have a simple model that could still provide useful predictions.

##### Cell cycle model overview

The cell cycle model is based on the conceptual "brake" model described by Lemmens and Lindqvist (Lemmens & Lindqvist, 2019). Briefly; once a cell is triggered to pass the restriction point, the cell is committed to division, and several brakes are kept in place to ensure orderly progression through the cell cycle. We implemented this conceptual model here into a mathematical model incorporating four brakes in the signaling network. A DNA licensing brake ensures that DNA licensing is at a sufficient level before replication starts (implemented through a brake on cyclin E/CDK2 activity while there is unlicensed DNA); a DNA replication brake ensures that DNA replication is finished before leaving S-phase (implemented through a brake on CDK1 activity while there is actively replicating DNA); a G2/M brake ensures that G2 is completed before mitosis starts (implemented through a brake keeping Wee1 active while G2 is ongoing); and the spindle assembly checkpoint ensures that the mitotic spindle is formed before cytokinesis (implemented through a brake on CDC20 activity while the spindle is not assembled). On this basic scaffold, additional mechanistic detail was added as described below.

#### Gene transcription

In the mitosis model, gene transcription was not specifically modelled to keep the initial test model small. In the cell cycle model, transcripts were modelled explicitly, as transcriptional regulation seemed to be necessary to explain the dynamics of various proteins. We assumed that transcription follows Hill kinetics with a fixed Hill coefficient of 4 and $K_d$ of 0.3. The rates of transcription were allowed to vary between genes for genes where we have measurements of the transcript or protein, and for other genes was set at the fixed rate of $1.1 \cdot 10^{-3}$, resulting in transcript synthesis in 15 minutes. Transcriptional inhibition (in the case of transcription of E2F7-regulated genes) is modelled by multiplying the transcription rate with one minus a second hill function, again with a $K_d$ of 0.3 but a Hill coefficient of 10 to allow the inhibiting transcription factor to strongly inhibit transcription when present at high levels in a cell.

#### Protein synthesis

We modeled protein synthesis with a separate molecular species representing the complex of actively translating messenger RNAs together with ribosomes, which are converted into a nascent full protein and a recovered messenger RNA at a fixed rate of $8.3 \cdot 10^{-4}$, resulting in protein synthesis in 20 minutes. We further assumed that the rate of protein synthesis decreases linearly with the amount of protein product present, with protein synthesis stopping entirely when a level of 1 is reached; implemented by decreasing the rate of translation-complex initiation by the sum of translating complexes plus full protein (in complex with other proteins if applicable). The main reason to include this product inhibition is to decouple the rates of synthesis and degradation; without this assumption, a protein-specific degradation rate is needed to balance production rates when the protein is at steady state. Such a protein-specific degradation and synthesis rate is typically unidentifiable given the measurements that were used here. One exception is made; namely for cyclin A – here we allow the protein level to reach 2 instead of 1. This is an ad-hoc choice; at steady state, half of the cyclin A will bind with either of the two CDKs in the model, so allowing cyclin A to reach 2 will allow cyclin A in complex with CDK2 to be comparable to the amount of cyclin E in complex with CDK2, such that we can keep the parameters of CDK2 activity the same for most reactions catalyzed by CDK2 in complex with either cyclin.

#### Proteasomal degradation

Many proteins in the models are subject to active degradation by proteasomes, which is regulated by E3 ubiquitin ligases. Protein degradation by proteasomes typically follows Michaelis-Menten kinetics (Luciani *et al*, 2005), as can also be seen in the mostly linear rather than exponential degradation of exogenous GFP-tagged cyclin B (Clute & Pines, 1999; Geley *et al*, 2001). The Michaelis-Menten constant

$K_M$ for degradation is somewhat arbitrary in the model, and set to 0.001 such that degradation is linear until the protein levels are low. For the catalytic rate constant $k_{cat}$ we used the fixed value of $1.6 \cdot 10^{-3}$; this combination of $K_M$ and $k_{cat}$ gives a near-complete degradation of a protein in 10 minutes, when the corresponding E3 ubiquitin ligase is fully active.

Two exceptions in degradation are cyclin E and p21. For cyclin E we included separate rate constants for the degradation of inactive cyclin E (slow, exponential decay) while retaining Michaelis-Menten-type degradation of active cyclin E (representing auto-phosphorylation and Fbw7-dependent proteasomal degradation, (Welcker *et al*, 2003)). P21 required an additional degradation term (on top of SKP2-dependent proteasomal degradation) to be able to describe variability between cells that have transient p21 expression and cells that have sustained p21 expression.

#### Protein activation/deactivation

For various proteins we explicitly modeled the inactive and active states; these states summarize one or more regulatory processes simultaneously. For example, cyclin E can be either in an inactive or active state. The active state represents binding to CDK2, as well as the absence of inhibitory proteins like p21 and p27. By not explicitly modeling all the possible configurations of protein complexes and phosphorylation states, we can keep the number of molecular species as low as possible, while still being able to represent contextual activation and inactivation.

#### Mitogenic signaling

The main focus of the model is to describe progression through the cell cycle rather than entry into the cell cycle. The extensive process of mitogenic signaling is therefore summarized into a single variable that increases over time. This variable could represent ongoing serum stimulation or an active oncogene. The mitogenic signal leads to cell cycle entry through three routes: first, stimulation of E2F1 expression (representing relief of repression by p130 (Smith *et al*, 1996) as well as the contribution of Myc (Leung *et al*, 2008)), second, activation of E2F1 (representing phosphorylation of Rb by cyclin D/CDK4, although this process is still not fully understood (Rubin *et al*, 2020)), and third, the inactivation of p27 (representing multiple regulatory steps (Vervoorts & Lüscher, 2008)).

#### DNA replication

DNA was modeled with four distinct molecular species, representing unlicensed DNA, licensed DNA that is primed for replication, DNA that is actively being replicated, and nascent replicated DNA. DNA needs to be licensed before cells can enter S-phase. The cell cycle brake on the extent of DNA licensing was modelled through an inhibition of CDK2 activity while there is unlicensed DNA (Nevis *et al*, 2009). Once DNA is licensed, active CDK2 along with the replisome can fire origins to convert the licensed DNA into actively replicating DNA. In the model, this process of origin firing is limited by the amount of active replicating DNA, which keeps the fraction of DNA that is replicating at any moment to a maximum of 10%. This represents the biological process that some origins are fired early while others are fired later. Replicating DNA will produce nascent replicated DNA at a steady rate. In the model, replicating DNA inhibits CDK1 to prevent S-phase exit while DNA replication is ongoing. Finally, replicating DNA can induce DNA damage, representing DNA damage caused by endogenous replication stress.

#### Mitosis

In the model, cells enter mitosis when cyclin B/CDK1 activity reaches the threshold of 0.5. At this point, the nuclear envelope is broken down, spindle components and the spindle assembly checkpoint are synthesized, and the spindle assembly is initiated. As long as spindle assembly is not completed, the spindle assembly checkpoint keeps CDC20 in an inactive state; when spindle assembly is finished the spindle assembly checkpoint is removed and CDC20 is allowed to degrade cyclin B and other cell cycle components. CDC20 also initiates chromatid separation (representing the release of separase due to degradation of securin by CDC20). Once the chromatids are separated, cytokinesis is initiated.

##### Cell division

Once the dynamic variable representing cytokinesis reaches a value of 1, the cell is removed from the simulation and two new cells are added. The level of all molecular species in the two daughter cells are set to be the same as the parent cell, with several exceptions: all four DNA species are divided by two, the species representing major cellular processes (including mitogenic signal, unassembled and assembled spindle, chromatid separation, cytokinesis and G2 delay) are all set to 0 if they were not yet 0 already, and the nuclear envelope is reset to 1. Variability in kinetic rate constants or species are then applied to the two daughter cells as described in the section on cell variability in the main text Materials and Methods.

#### ODE integration

To integrate each ODE system, we used the CVODE algorithm of the SUNDIALS package (Hindmarsh *et al*, 2005; Gardner *et al*, 2022), using the Backward Differentiation Formulas as the ODE systems tend to be stiff. To further increase the efficiency of the ODE integration for this already efficient solver, we incorporated three computational optimizations.

First, we used a custom linear solver during the Newton-Raphson iterations. For biochemical reaction networks, the Jacobian of the ODE system is usually quite sparse, since one molecular species typically interacts with only a few other molecular species. However, in our models the overall size of the ODE system is still relatively modest, and we found that the overhead of dedicated sparse solvers is too large, resulting in slower simulations than using a dense solver. On the other hand, the built-in dense solver in the SUNDIALS package does not make optimal use of CPU vectorization. We therefore used the PartialPivLU solver of the Eigen C++ library which does have optimized vectorization, with two modifications. First, we skip over zero entries where possible, which is beneficial here given the relative sparse Jacobians; and second, we disabled the blocking feature, as the ODE systems are sufficiently small to do the LU-decompositions in an unblocked way still efficiently in the CPU caches.

Second, we automatically generate C++ code for evaluating the time derivatives and Jacobian matrices from the rate law equations specified in the model, and automatically compile this to a dynamic library that can be used by the ODE integrator.

Third, several optimized functions for evaluating the rate laws are included, especially for fast evaluation of hill functions with specific integer coefficients.

Together, these three optimizations resulted in a 4-fold improvement over the initial implementation, and a 15-fold performance gain for simulating a single cell compared to simulation in COPASI (Hoops *et al*, 2006) with identical tolerances.

The parallel tempering inference scheme used for parameter inference provides some opportunity for parallelization, but a dedicated compute server typically has more computing cores available than the optimal number of parallel chains to be used. We therefore parallelized the inference not just over the parallel sampling chains, but also over the cells. While simulating a population of cells, we integrate the ODE system for all cells separately and store the Nordsieck arrays of the solver at each ODE integration step in memory, so that any intermediate time point can be interpolated at the end of the simulation. This storing of integration points is needed when cell trajectories are synchronized to a simulation event, such as the nuclear envelope breakdown, since the time point of the measurement relative to the simulation event will not be known in advance in such cases. This parallelization scheme prohibits communication between cells, but greatly speeds up the simulation when many computing cores are available, as all cells can then be simulated independently.

#### Choice of inference parameters

Parameter inference in combination with simulating populations of cells is challenging, as the computational cost scales cubically in the number of dynamic variables in the model, and (in the worst case) exponentially in the number of inference parameters, multiplied by the number of cells and number of datasets. The most promising route to keep the computation manageable therefore seems to be to keep the number of inference parameters as low as possible. To that end, we made several simplifying assumptions in the construction of the model (as described above). Furthermore, we fixed as many parameters as possible to specific values based on assumptions or literature, and we left only the most important model parameters free for inference, described in more detail below.

##### Mitosis model

For the mitosis model, seventeen or sixteen inference variables were estimated from the two sets of measurements respectively. This includes six inference variables for the model parameters that vary between cells (the means and standard deviations of FOXM1 synthesis, mitotic entry rate and spindle assembly rate), seven inference variables for model parameters that do not vary between cells, an inference variable for the relative standard deviation of the error term, and either three or two additional variables specific for each dataset (a CDK1 sensor activation-, deactivation- and dilution-parameter for the CDK1 sensor measurements, and an offset and scale parameter for the eYFP-tagged cyclin B measurements).

##### Cell cycle model

For the cell cycle model, we used between nine and twenty inference variables depending on the dataset. Specifically, we selected only the mean rates and the variabilities of the five main cellular processes as unknown variables, plus the rates of transcription of the specific gene products that are being measured in the particular dataset that is being fitted, an entry time if necessary to generate variable cells at the start of the trajectory, and the standard deviation of the error term. For instance, the dataset of Eward et al contained measurements of cyclin E, A and B; when fitting this dataset we used 15 variables: five means and five standard deviations for the five main cellular processes, three transcription rates (for cyclin E, A and B), the standard deviation of the error term, and an entry time.

The five main cellular processes were, as described in the main text as well: 1) the rate of mitogenic signal accumulation, 2) the rate of DNA licensing, 3) the rate of DNA replication, 4) the rate of endogenous replication stress-induced DNA damage, and 5) the rate at which cells progress through G2. When a particular cellular process rate had no influence on the measurements, variability was not included as inference parameter. For example, the measurements of Cappell et al correspond only to G1 and S phase, so variabilities in endogenous replication-stress-induced DNA damage and G2 delay were not included.

#### Parameter inference

Parameter inference was done with Bayesian statistics, using parallel-tempered Markov chain Monte Carlo sampling (Earl & Deem, 2005) with a custom, locally adaptive proposal distribution for the mutate moves.

The parallel chains were set up to sample from the fractional posteriors

$$P_\eta(\boldsymbol{\theta}|\mathbf{y}) \sim P(\mathbf{y}|\boldsymbol{\theta})^\eta P(\boldsymbol{\theta})$$

where the learning rate $\eta$ was distributed between the chains according to a power law schedule. The first chain (i.e., the chain with the lowest $\eta$, and consequently highest temperature) has $\eta$ equal to zero so that this chain samples directly from the prior. The last chain has $\eta$ equal to one, so that it samples from the full posterior. The $\eta$ of the intermediate chains was set to $\eta_i = (i-1)/(n_c-1)^\gamma$, where $n_c$ is the number of chains. The number of chains and power γ were chosen such that there were no bottlenecks in swapping of chain pairs (that is, all pairs had to have an acceptance rate of at least 0.1 during swap moves), as well as providing a sufficient range of learning rates for the posterior calibration process. Twenty-four parallel chains with a $\gamma$ of 3 for the mitosis model and 4 for the cell cycle model appeared to be sufficient.

To select between swap moves and mutates moves in the parallel tempering, we used the deterministic even-odd scheme (Syed *et al*, 2022; Okabe *et al*, 2001). That is, we deterministically alternate between a swap move followed by a number of mutate moves. For the mitosis model, we used 1 mutate move for every swap, and for the cell cycle model we used 2 mutates moves for every swap. In each swap move, we alternate between attempting a swap for every even and odd pair. That is, in the first swap move we attempt to swap chains 1 and 2, 3 and 4, 5 and 6, etc; and in the following swap move we attempt to swap chains 2 and 3, 4 and 5 and the last and first chain.

As proposal distribution for mutate moves in each parallel chain, we used a random walk proposal using a Gaussian mixture distribution adapted to the sample chain history. In every run, we divide the sampling process into four sections. In the first section, we sample with a single diagonal Gaussian distribution with variances based on the prior distribution. In the second and third section, we sample with a Gaussian mixture fitted to the samples obtained from the previous section. In each of these three sections, the scales of the Gaussians are continuously adapted to obtain a good acceptance rate. Finally, in the fourth section, the Gaussian distribution as well as the adaptive scaling is fixed to those obtained at the end of the third section. Only the samples of the fourth section are used as the final samples for downstream analysis. The first three sections each correspond to $1/6^{th}$ of the total samples, and the final fourth section corresponds to $3/6^{th}$ of the total samples.

The Gaussian mixture distributions are fitted to the historical samples using expectation-maximization. The historical samples typically have autocorrelation, and the chain typically has not converged yet when the Gaussian mixture is fitted. We therefore estimate the covariance during expectation-maximization with shrinkage; using the orthogonally invariant minimax estimator described in (Dey & Srinivasan, 1985). That is, we do an Eigen decomposition of the empirical covariance matrix, and scale the eigenvalues according to

$$v_i^* = v_i \frac{n}{n+d+1-2i}$$

where $v_i$ is the $i$th eigenvalue sorted from high to low, $n$ is the number of samples and $d$ is the number of dimensions. For $n$, we used the sum of the weights of the samples based on the responsibilities of each mixture component, adjusted by a factor based on the effective sample size, to account for autocorrelation in the samples. When this value $n$ is less than the number of variables, we scale the first n eigenvalues according to the formula above, and set the remaining eigenvalues to zero. We attempt to fit a Gaussian mixture with increasing number of components according to a Fibonacci sequence, until the number of components multiplied by half the number of variables plus one exceeds the number of available effective samples. Among these Gaussian mixtures with increasing number of components, the mixture with the lowest AIC is selected.

During sampling, one component of the Gaussian mixture is selected, using weighted sampling according to the responsibilities of the mixture at the current position of the chain, and the covariance of that component is used to generate a move in the random walk. During the sampling sections where the Gaussian mixture scale is adapted, we track how often a mutate move from each component of the mixture is accepted, using an exponential moving average. If the acceptance rate becomes too low or too high, the scale of that component is decreased or increased respectively, aiming for an acceptance rate of 0.234 when the number of dimensions is 4 or higher, or somewhat higher values for lower dimensions (Roberts & Rosenthal, 2001).

### Supplementary results

#### Supplementary Result 1 – Computational efficiency

As described in the Supplementary Methods, we implemented four optimizations to increase the computational efficiency of simulating the population of cells; three optimizations to improve individual ODE integrations, and the fourth optimization to parallelize the simulation efficiently. To compare the effect of the three optimizations of the ODE integration, we simulated the mitosis model 5,000 times, with parameter values taken from Monte Carlo samples from all temperatures of the fit to the measurements of Gavet & Pines, on a single core. Without any optimizations, the C++ implementation using default CVODE solver achieved 12 ODE integrations/second. By generating C++ code for evaluating the rate law and Jacobian matrix evaluations, this increased to 39 ODE integrations/seconds. Using optimizations for specific functions such as the hill function with integer coefficients this further increased to 50 ODE integrations/second, and using the optimized linear solver leveraging code vectorization as well skipping over zero entries where possible, this finally resulted in 56 ODE integrations/second. For reference, CopasiSE (version 4.39) achieved 3.6 ODE integrations/second on average on the same sets of parameter values. Together, this represents a factor 15 improvement over an available tool for ODE integration of SBML models, and a factor 4.7 over an initial implementation with available libraries.

To compare the efficiency of several sampling algorithms, we focused on the fit of the mitosis model to the measurements of Akopyan et al (described in the main text), using seven instead of sixteen inference parameters to keep the computation time manageable when using alternative sampling algorithms. We compared three algorithms. First, we used a global covariance matrix adapted to the sample history (Haario *et al*, 2001), adaptively scaled to give an acceptance rate of 0.234; second, we used the autoblocking strategy described in (Turek *et al*, 2017) (this was the previous default sampling algorithm in the BCM software package); and third, the adaptive shrunken Gaussian mixture described in the present manuscript. Each run used the same number of likelihood evaluations (9.6 million likelihood evaluations, to produce 12,000 posterior samples of which the first half was used for adaptation and discarded as burn-in). Supplementary Figure 1A and B show trace and autocorrelation plots of these three algorithms, clearly showing reduced autocorrelation using the mixture proposal. The mixture proposal is adapted to the local shape of the target probability distribution, with two example bivariate scatter plots illustrating the fit (Supplementary Figure 1C). Given the 6,000 posterior samples (after burn-in discarding), the global covariance proposal produced 185 effective samples (calculated as the minimum effective sample size over the marginals of the seven inference parameters), the autoblocking algorithm produced 224 effective samples and the mixture proposal produced 740 effective samples; giving a factor 4 and 3.3 improvement respectively. Multiplied with the 4.7-fold improvement of ODE integration efficiency described in the previous paragraph, this amounts to a 15-fold improvement over the initial implementation based on existing tools (the previous of BCM, standard CVODE integration with its reference linear solver, and reference rate law evaluation functions).

#### Supplementary Result 2 - Posterior calibration of cell cycle model

For the calibration of the posterior of the cell cycle model, we used two sets of measurements of the duration of S-phase (Burgess *et al*, 2012; Grant *et al*, 2018), which provided measurements for HeLa cells and U2OS cells respectively. There were three inference datasets that constrained the duration of S-phase: the data from Akopyan et al (U2OS cells), Cappell et al (HeLa cells), and Westendorp et al (HeLa cells); the measurements in these datasets spanned full S-phases. The other datasets did not necessarily cover full S-phases, or we did not have S-phase duration measurements of the relevant cell line. We let the cell cycle model predict the duration of S-phase, defined as the period of time between the point at which the dynamic variable representing replicating DNA first rose above $10^{-4}$, until the point at which the dynamic variable representing replicated DNA reached 1.95. The predictions based on Cappell et al and Westendorp et al were compared to the S-phase duration measurements of Burgess et al, and the predictions based on Akopyan et al were compared to the S-phase duration measurements of Grant et al (given those were in the same cell line).

The calibration result is shown in Supplementary Figure 6. The predictions of the S-phase duration of HeLa cells based on the data of Cappell et al are quite short, with many simulated cells progressing through S-phase within 2-3 hours, which seems unreasonably fast. The measurements of Cappell et al indicate a doubling of EdU signal within ~5 hours on average. Given that the model apparently needs some variability in S-phase entry, it needs fast progression through S-phase for some cells to reproduce the average EdU signal trace. It seems likely that the model is overestimating the variability in S-phase entry (leading to short S-phase durations to reproduce the average behavior), or the EdU signals may miss the earliest and latest stages of DNA replication. Either way a fairly strong correction is needed here (a learning rate of 0.14).

The predicted S-phase durations based on the fit to Akopyan et al are also clearly lower than what is observed in the measurements of Grant et al. It seems likely that averaged measurements of cyclin A and cyclin B do not provide sufficient information for an accurate estimation of S-phase duration, and that the predictions are overly confident in this case, again giving a fairly strong correction (learning rate of 0.1).

The predicted S-phase durations based on the fit to Westendorp et al are in line with measurements of Burgess et al. This is mainly due to the relatively low amount of information regarding S-phase durations obtained from western blots of E2F1 and E2F7 in synchronized cells at only a few time points, leading to a large uncertainty in the predictions of the duration of S-phase. The predicted error bars of the S-phase durations are indeed sufficiently wide such that the measurements in the external validation set fall within the 90% confidence intervals. Based on this comparison of these predictions from the fit to the data of Westendorp et al, a somewhat more modest calibration would be needed (learning rate of 0.3).

To err on the side of caution, we selected the minimum learning rate across these three comparisons. That is, we selected the optimal learning rate for each comparison separately using the method described in the main text, and then selected the lowest learning rate among the comparisons.

### Supplementary Figures

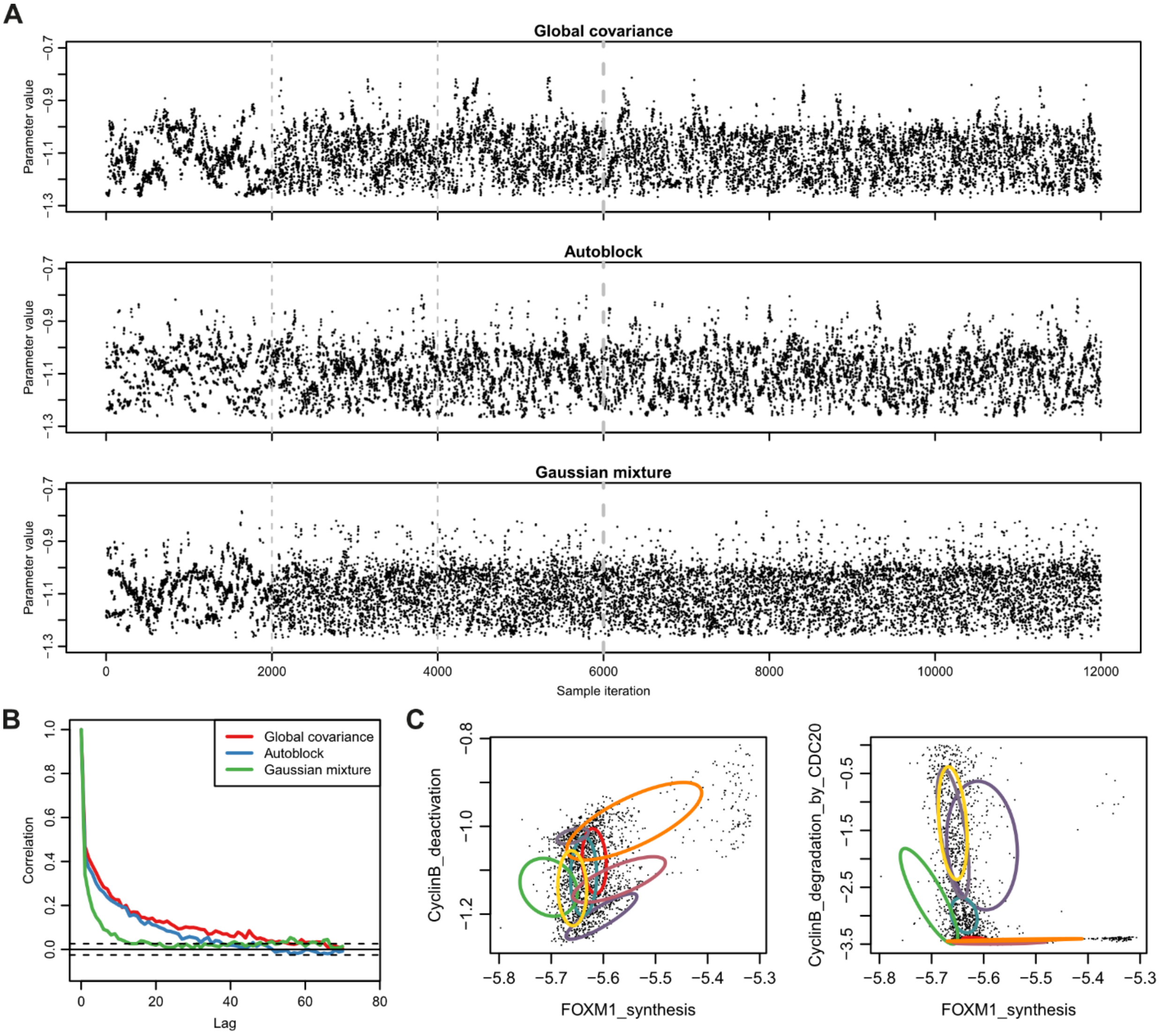


**Supplementary Figure 1: Sampler efficiency comparison. (A)** Trace plots of three inference runs: top - a Gaussian proposal distribution based on a global covariance matrix, middle- automated variable blocking based on Turek et al.'s algorithm, and bottom - the adaptive shrunken Gaussian mixture described in the present manuscript. Grey lines indicate the points in the chain where adaptation is applied (see Supplementary Methods). Traces are from the cyclin B deactivation rate parameter. **(B)** Autocorrelation plots for the cyclin B deactivation rate parameter (corresponding to the traces shown in A). **(C)** Visualization of the fitted Gaussian mixture proposal distribution, projected on two 2D scatter plots of three parameters. Each colored ellipse depicts one of the components of the Gaussian mixture distribution.

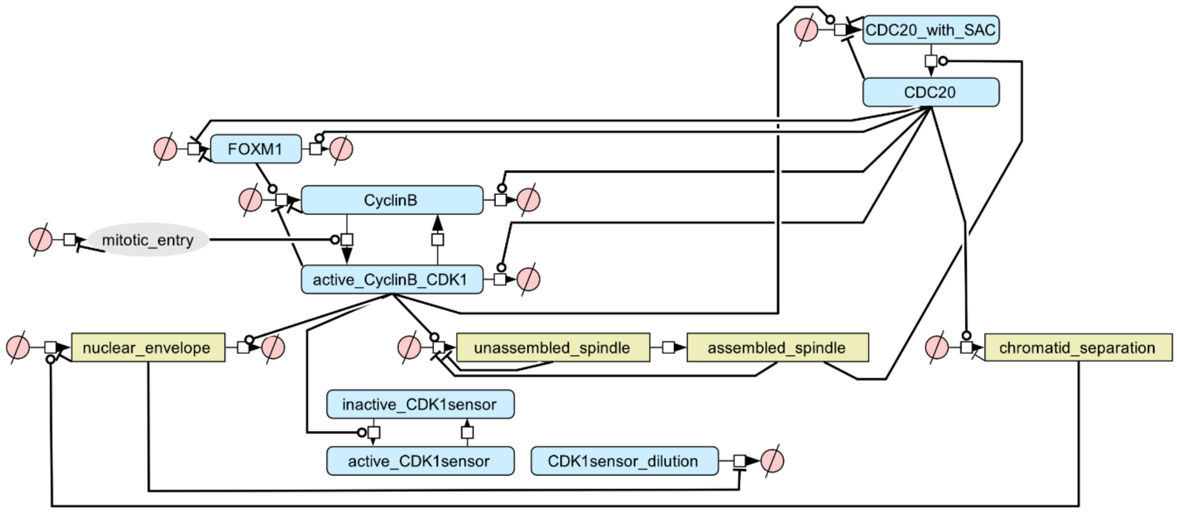


**Supplementary Figure 2: misspecified model used in the validation of the calibration process.** The model is identical to the main mitosis model, but with the positive feedback through CDC25C removed. When CDK1 activity increases above a certain threshold, it still activates the processes of nuclear envelope breakdown and mitotic spindle assembly. The main difference with the main mitosis model is that the feedback loop through CDC25C is not present to make the increase in CDK1 activity more switch-like.

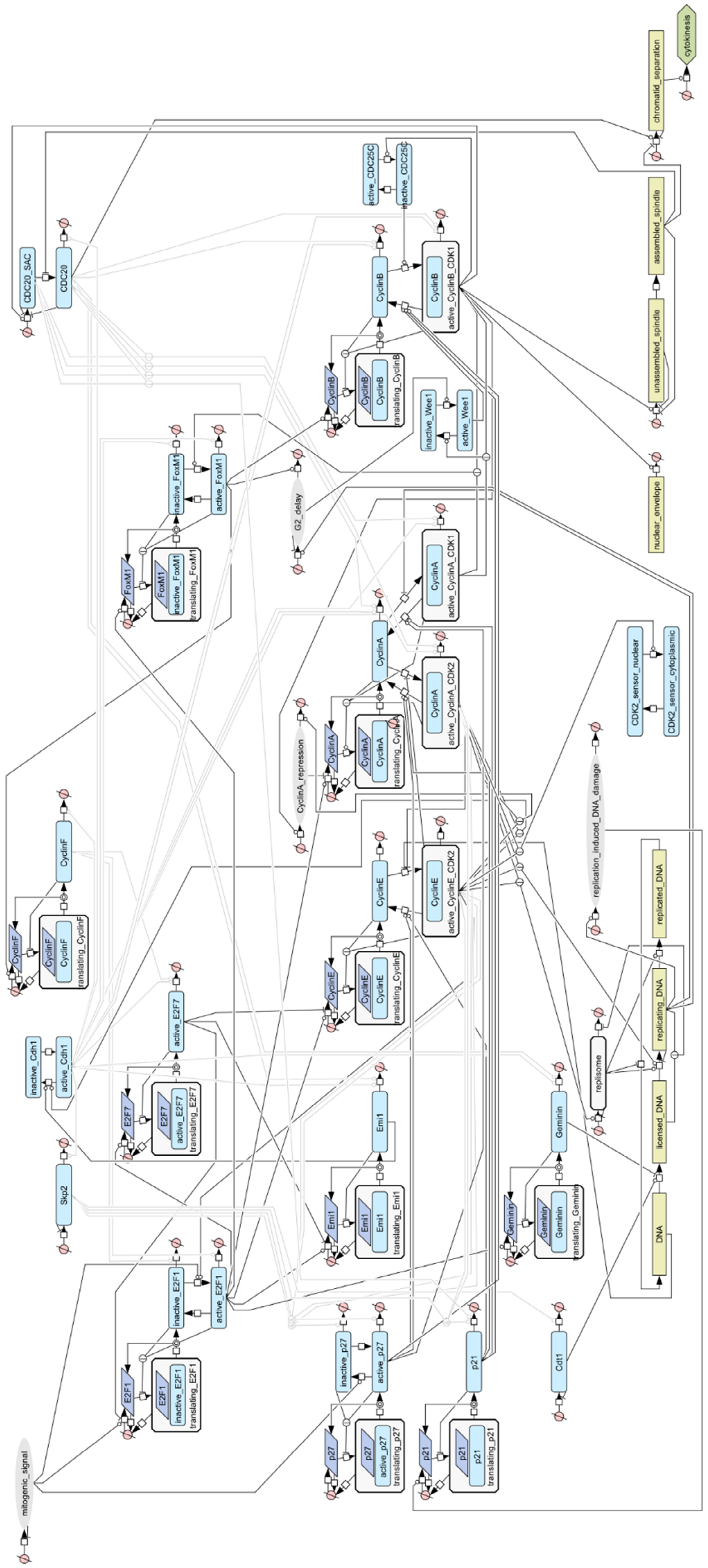


**Supplementary Figure 3: Larger view of the cell cycle model diagram**

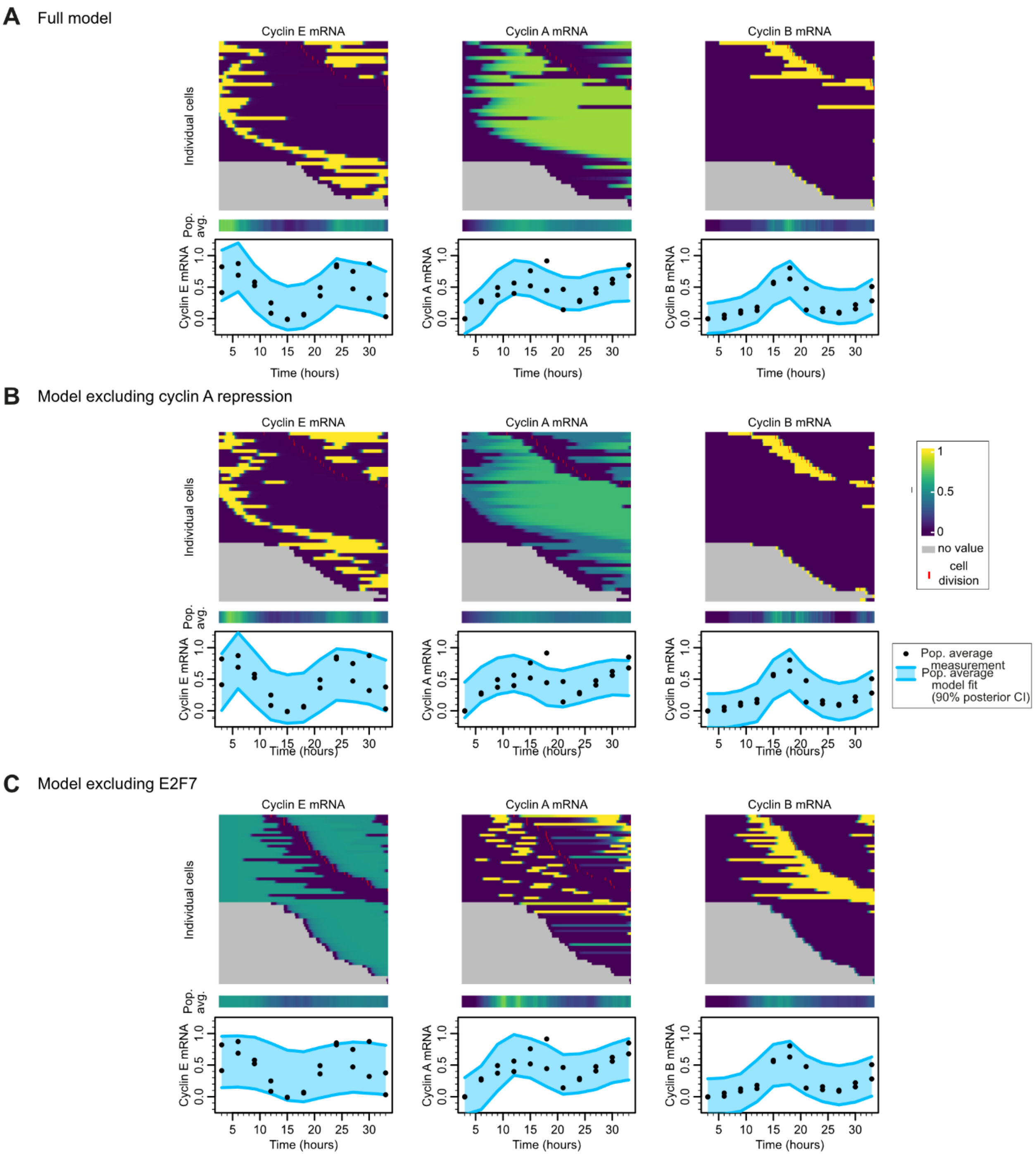


**Supplementary Figure 4: Trajectories and posterior predictive plots of the three model versions.** Related to main Figure 5. Each model was fit simultaneously on all three cyclin mRNA measurements; only E and A were shown in the main figure for brevity. **(A)** Trajectories and posterior predictive of the full cell cycle model. **(B)** Trajectories and posterior predictive of the cell cycle model excluding the cyclin A repression mechanism. **(C)** Trajectories and posterior predictive of the cell cycle model excluding E2F7.

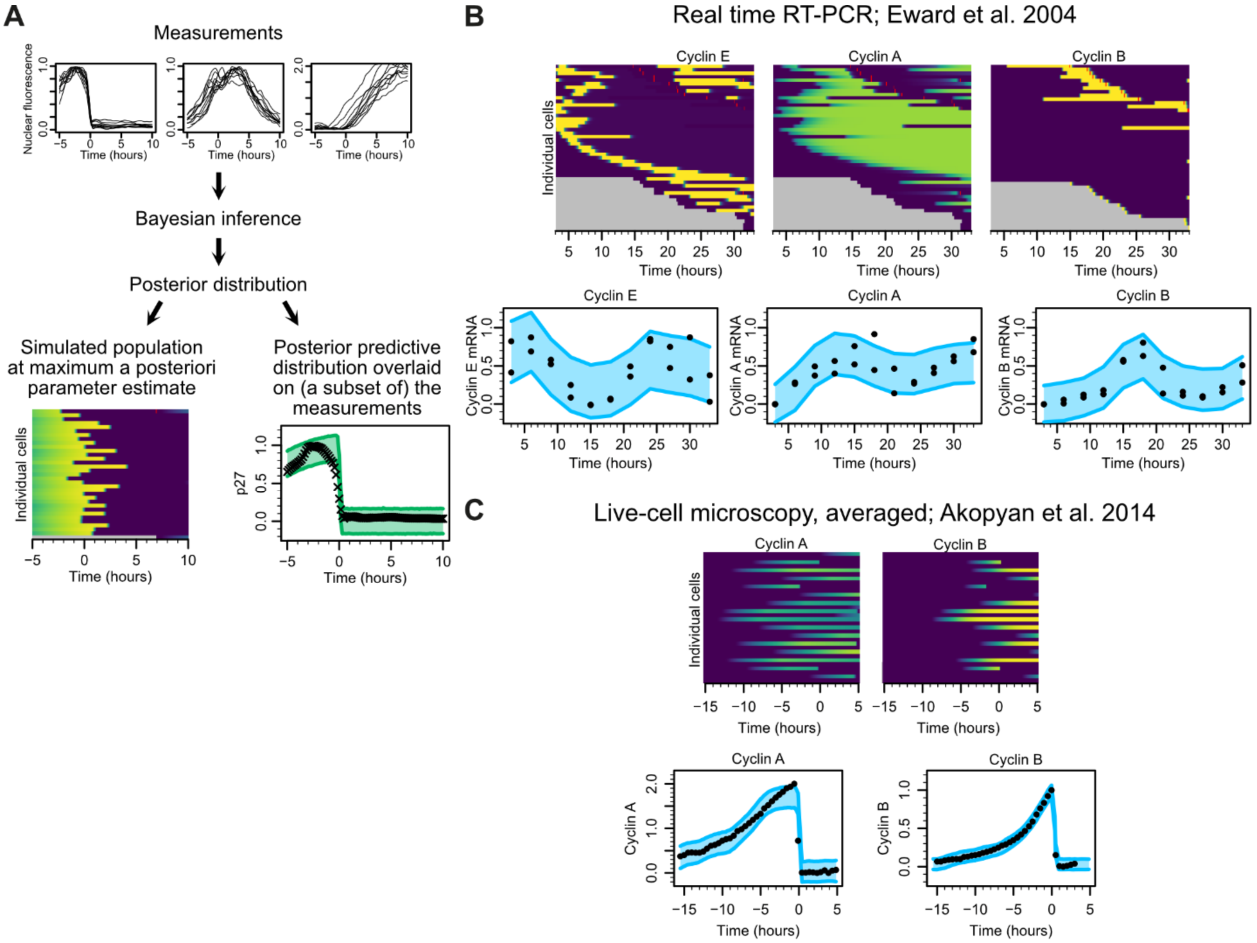


**Supplementary Figure 5: Cell cycle model simulations and fits to two additional datasets (extension of main Figure 6).** See legend of main Figure 6. **(A)** Illustration of how the trajectories and model fit figures are constructed. **(B)** Measurements of cyclin E, A and B mRNA levels measured by real-time RT-PCR of synchronized cells. Data was presented as generations in the publication of Eward et al Figure 2); this was converted to hours based on the cell cycle phase distributions (Figure 1 of Eward et al.). Two of the three fits were also shown in main Figure 5. **(C)**. Measurements of endogenous GFP-tagged cyclin A and B from Akopyan et al. The cyclin B measurements were also used with the mitosis model in Figure 2.

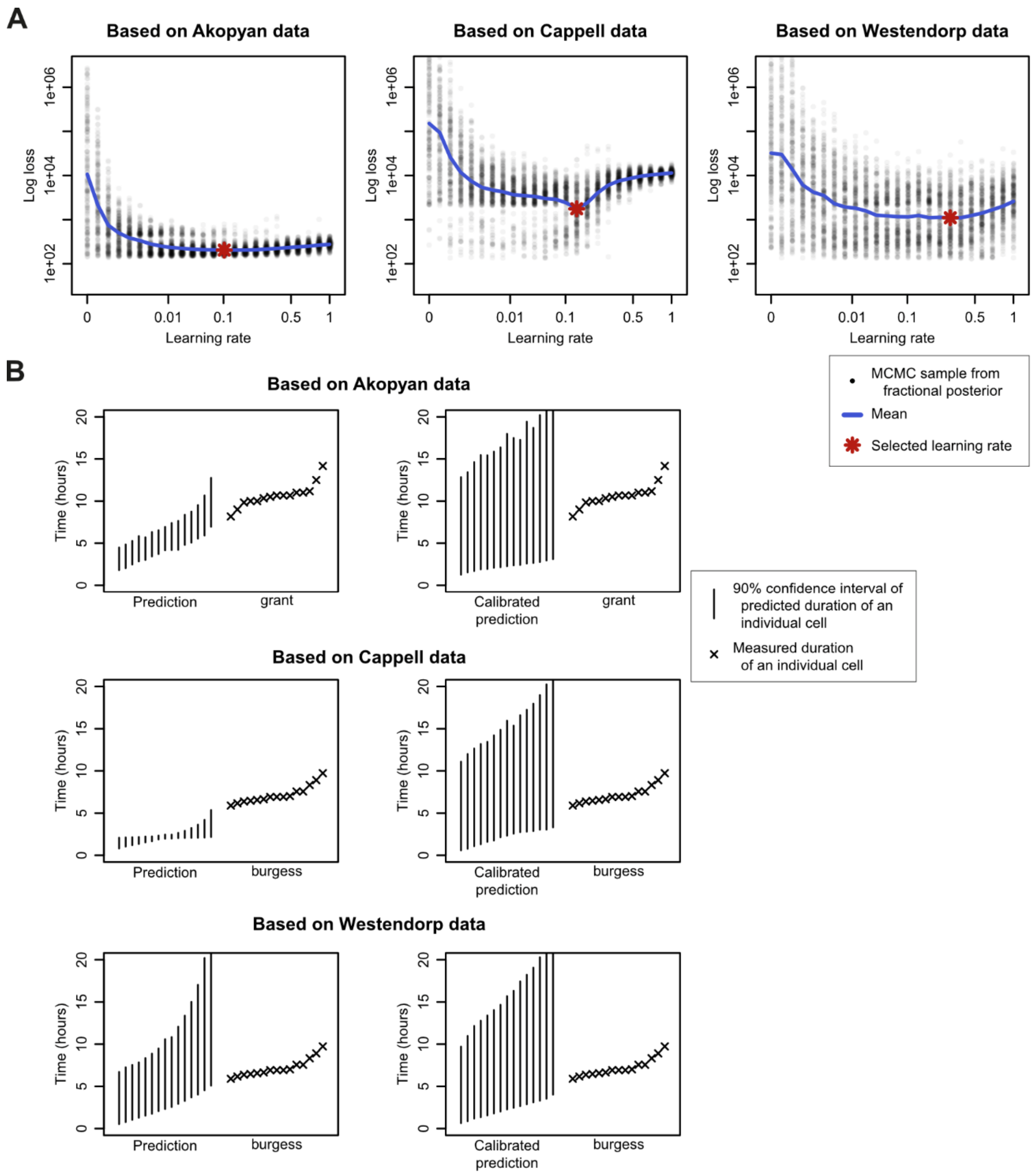


**Supplementary Figure 6. Posterior calibration of cell cycle model. (A)** Posterior calibration curves for the fit on the measurements of Akopyan, Cappell and Westendorp respectively. (see Methods and the legend of main Figure 3). **(B)** Comparison of predicted S-phase durations based on regular posterior (left column) and on the calibrated posterior (right column).

#### Supplementary References

Burgess A, Lorca T & Castro A (2012) Quantitative Live Imaging of Endogenous DNA Replication in Mammalian Cells. *PLoS ONE* 7: e45726

Clute P & Pines J (1999) Temporal and spatial control of cyclin B1 destruction in metaphase. *Nat Cell Biol* 1: 82–87

Dey DK & Srinivasan C (1985) Estimation of a Covariance Matrix under Stein's Loss. *Ann Statist* 13

Earl DJ & Deem MW (2005) Parallel tempering: Theory, applications, and new perspectives. *Phys Chem Chem Phys* 7: 3910

Gardner DJ, Reynolds DR, Woodward CS & Balos CJ (2022) Enabling New Flexibility in the SUNDIALS Suite of Nonlinear and Differential/Algebraic Equation Solvers. *ACM Trans Math Softw* 48: 1–24

Geley S, Kramer E, Gieffers C, Gannon J, Peters J-M & Hunt T (2001) Anaphase-Promoting Complex/Cyclosome–Dependent Proteolysis of Human Cyclin a Starts at the Beginning of Mitosis and Is Not Subject to the Spindle Assembly Checkpoint. *Journal of Cell Biology* 153: 137–148

Grant GD, Kedziora KM, Limas JC, Cook JG & Purvis JE (2018) Accurate delineation of cell cycle phase transitions in living cells with PIP-FUCCI. *Cell Cycle* 17: 2496–2516

Haario H, Saksman E & Tamminen J (2001) An Adaptive Metropolis Algorithm. *Bernoulli* 7: 223

Hindmarsh AC, Brown PN, Grant KE, Lee SL, Serban R, Shumaker DE & Woodward CS (2005) SUNDIALS: Suite of nonlinear and differential/algebraic equation solvers. *ACM Trans Math Softw* 31: 363–396

Hoops S, Sahle S, Gauges R, Lee C, Pahle J, Simus N, Singhal M, Xu L, Mendes P & Kummer U (2006) COPASI—a COmplex PAthway SImulator. *Bioinformatics* 22: 3067–3074

Lemmens B & Lindqvist A (2019) DNA replication and mitotic entry: A brake model for cell cycle progression. *Journal of Cell Biology* 218: 3892–3902

Leung JY, Ehmann GL, Giangrande PH & Nevins JR (2008) A role for Myc in facilitating transcription activation by E2F1. *Oncogene* 27: 4172–4179

Luciani F, Keşmir C, Mishto M, Or-Guil M & de Boer RJ (2005) A Mathematical Model of Protein Degradation by the Proteasome. *Biophysical Journal* 88: 2422–2432

Nevis KR, Cordeiro-Stone M & Cook JG (2009) Origin licensing and p53 status regulate Cdk2 activity during G1. *Cell Cycle* 8: 1952–1963

Okabe T, Kawata M, Okamoto Y & Mikami M (2001) Replica-exchange Monte Carlo method for the isobaric–isothermal ensemble. *Chemical Physics Letters* 335: 435–439

Roberts GO & Rosenthal JS (2001) Optimal scaling for various Metropolis-Hastings algorithms. *Statist Sci* 16

Rubin SM, Sage J & Skotheim JM (2020) Integrating Old and New Paradigms of G1/S Control. *Molecular Cell* 80: 183–192

Smith EJ, Leone G, DeGregori J, Jakoi L & Nevins JR (1996) The accumulation of an E2F-p130 transcriptional repressor distinguishes a G0 cell state from a G1 cell state. *Mol Cell Biol* 16: 6965–6976

Syed S, Bouchard-Côté A, Deligiannidis G & Doucet A (2022) Non-Reversible Parallel Tempering: a Scalable Highly Parallel MCMC Scheme. *J R Stat Soc Series B* 84: 321–350

Turek D, de Valpine P, Paciorek CJ & Anderson-Bergman C (2017) Automated Parameter Blocking for Efficient Markov Chain Monte Carlo Sampling. *Bayesian Anal* 12

Vervoorts J & Lüscher B (2008) Post-translational regulation of the tumor suppressor p27KIP1. *Cell Mol Life Sci* 65: 3255–3264

Welcker M, Singer J, Loeb KR, Grim J, Bloecher A, Gurien-West M, Clurman BE & Roberts JM (2003) Multisite Phosphorylation by Cdk2 and GSK3 Controls Cyclin E Degradation. *Molecular Cell* 12: 381–392
